# Distributed representations for cognitive control in frontal medial cortex

**DOI:** 10.1101/2023.12.12.571242

**Authors:** Thomas R. Colin, Iris Ikink, Clay B. Holroyd

**Affiliations:** Gent University; Ghent University

**Author notes:** corresponding author Department of Experimental Psychology Ghent University Ghent, 9000, Belgium.

## Abstract

In natural and artificial neural networks, modularity and distributed structure afford complementary but competing benefits. The former allows for hierarchical representations that can flexibly recombine modules to address novel problems, whereas the latter affords better generalization. Here we investigate these competing demands in the context of sequential behavior. First, we explore this by comparing the properties of several recurrent neural network models. We find that explicit hierarchical structure fails to provide an advantage for generalization above a “flat” model that does not incorporate hierarchical structure. However, hierarchy appears to facilitate cognitive control processes that support non-routine behaviors and behaviors that are carried out under computational stress. Second, we compare these models against functional magnetic resonance imaging (fMRI) data using representational similarity analysis. We find that a model that incorporates so-called wiring costs in the cost function, which produces a hierarchically-organized gradient of representational structure across the hidden layer of the neural network, best accounts for fMRI data collected from human participants in a previous study (Holroyd et al., 2018). The results reveal that the anterior cingulate cortex (ACC) encodes distributed representations of sequential task context along a rostro-caudal gradient of abstraction: rostral ACC encodes relatively abstract and temporally extended patterns of activity compared to those encoded by caudal ACC. These results provide insight into the role of ACC in motivation and cognitive control.

---

A perennial debate in cognitive science concerns the form of the computational representations that underpin cognition (Gardner, 1985; Hanson & Burr, 1990; Bowers, 2017; Garnelo & Shanahan, 2019). This debate has centered on the relative strengths of two complementary approaches to solving computational problems. On the one hand, it has been argued that cognition is mediated by discrete, symbolic codes, a perspective precipitated by the advent of the digital computer. This approach affords the flexible recombination of elementary units into modular and/or hierarchical structures, which is a hallmark of linguistic codes (Chomsky, 1957). By contrast, the alternative view holds that cognition is encoded in distributed, multivariate representations, an idea inspired by brain-like computations carried out by artificial neural networks (Rogers & McClelland, 2014) and their modern incarnation, deep learning algorithms (Schmidhuber, 2015). This approach trades the compositional flexibility associated with the symbolic approach for other computational benefits; in particular, an unmatched ability to learn representations for diverse and challenging tasks, and to generalize those to novel situations. A similar tension is also apparent in hierarchical reinforcement learning (RL), an influential theoretical framework that makes use of discrete, combinable behavioral components (termed “options”, Sutton et al., 1999), where more recent algorithms have leveraged distributed representations (see e.g. Bacon et al., 2017, Vezhnevets et al. 2017, Haarnoja et al., 2018). It is not clear how these seemingly diverging demands of distributed representations vs. hierarchical organization can best be reconciled.

Although this debate continues to echo in contemporary contexts (e.g., Lake et al., 2017), it bears noting that actual brains, which act in the crucible of the real world, must have already found an appropriate balance between the demands of hierarchy and distributed representations (Richards et al., 2019, Marblestone et al., 2016). This balance is perhaps most evident in frontal cortex, which encodes high-dimensional representations of task execution across ensembles of neurons, but which is also believed to implement cognitive control processes that operate at hierarchically-high levels of processing (Cohen, 2017). Further, these representations are proposed to be physically encoded along the rostral-caudal extent of frontal cortex in hierarchically-organized modules (Koechlin et al., 2003, Badre & Nee, 2018). Computational studies have yet to address fully how these tradeoffs are mediated by frontal systems in real brains, especially in humans, where the neural implementation of cognitive control processes might be expected to be especially complex.

In the present article, we address this question by extending a previously-developed recurrent neural network (RNN) model of action sequence production proposed to engage the control processes of the anterior cingulate cortex or ACC (Shahnazian & Holroyd, 2018). This model utilized distributed representations to predict action sequences, in line with previous treatments of this problem (Botvinick & Plaut, 2004, Cooper et al., 2014), but in contrast with recent proposals of ACC function (Holroyd & Yeung, 2012; Holroyd & McClure, 2015; Holroyd & Verguts, 2021), the model did not incorporate hierarchy into these representations. Accordingly, here we incorporated symbolic-like goal units representing goals and subgoals into the RNN model, conjecturing that these units would support goal-directed temporal patterns that enable flexible behavior. We first investigated the computational properties of the model (study 1) and then tested the neural viability of the model using an existing functional magnetic resonance imaging (fMRI) data set (study 2).

In particular, study 1 extended previous work that used RNNs to simulate action sequences (Botvinick & Plaut, 2004, Cooper et al., 2014, Holroyd et al, 2018) by integrating symbolic-like “goal” units into a standard (i.e., “flat”) RNN. We tested the model using a so-called *kitchen environment* expressly designed to encourage the development of representations that generalize across stimulus-response associations and temporally extended actions, while also affording modular composition of action patterns. Crucially, this hybrid approach allowed for investigating the interface between hierarchically-organized neural modules and their influence on sequence production mediated by distributed representations across the model’s internal “hidden” units. In so doing, the model provided a means to explore how high-level cognitive control processes can affect action-related representations carried out by other systems. For example, we examined whether enhanced activation of hierarchically high goal units (as proposed in classic models of cognitive control, e.g., Cohen et al., 1990) can appropriately bias action representations during the execution of novel action sequences.

In study 2, we used representational similarity analysis (RSA) to look for second-order similarities between representations encoded by the RNN when trained on a simplified version of the kitchen environment, and neural representations in humans carrying out the same task. To our knowledge this constitutes one of very few investigations of PFC using model-based RSA to date (Freund et al., 2021), aside from our own prior work (Holroyd et al., 2018) and work by Cross et al. (2021). Holroyd et al. (2018) revealed second-order similarities between the ‘flat’ representations encoded by the hidden units of the model and the representations encoded by anterior cingulate cortex (ACC: Shahnazian & Holroyd, 2018), a frontal brain region associated with monitoring hierarchical action sequences for the purpose of planning (Holroyd & Verguts, 2021). Given previous theoretical work suggesting that ACC is hierarchically organized, with relatively rostral parts of it concerned with higher levels of task execution (Holroyd & McClure, 2015), we expected that the hierarchical implementation of this RNN would reflect this rostral-caudal organization along ACC. This work provides a point of departure for studying the computational representations of cognitive control in ACC and other frontal systems.

## Study 1: Modeling

An early debate in cognitive psychology centered on whether explicitly hierarchical representations are necessary to effect goal-directed behavior. Lashley (1951) accounted for the limitations of associationist descriptions of behaviors using schemas, which gave rise to a literature emphasizing the hierarchical structure of behavior (Miller et al. 1960, Schank & Abelson 1977, Grafman 1995). However, in contrast with Lashley’s seminal work, Botvinick and Plaut (2004) utilized an RNN model of sequential decision making to demonstrate that explicit hierarchy is not necessary for neurocognitive systems to learn and carry out structured behaviors. In response, Cooper and colleagues (2014) demonstrated that augmenting RNNs with goal units can better explain observations of sequential behaviors by facilitating the application of cognitive control. For example, the goal units enabled flexible control at the level of previously learned subroutines, and with appropriate training, seemed to enable generalization to a wide range of starting states. Although classic computational models have shown that networks that incorporate goal units can effectively simulate the effect of top-down control signals mediated by prefrontal cortex over other neural systems responsible for carrying out automatic or overlearned behaviors (Miller & Cohen, 2001), the role of these units in hierarchically-organized, goal-directed sequential behaviors is still not well understood.

In study 1, we pursued this line of research further by implementing two RNNs – one with and one without goal units – and then subjecting them to a series of tests to evaluate their functional properties. For this purpose, we utilized a richer environment than that used by Botvinick and Plaut (2004) and Cooper et al. (2014), in order to provide the model with more opportunities to exploit statistical regularities in the task. We investigated whether the incorporation of goal units representing currently active goals and sub-goals imposed hierarchical structure onto the network representations, and whether these provided an efficient means for the application of top-down control. We also intervened on network activity in several ways that simulated external perturbations of the system related to distraction and fluctuations of control.

### The kitchen environment

The task, which was inspired by similar environments used by Botvinick and Plaut (2004) and Cooper et al. (2014), consisted of making tea or coffee: given sensory input and using the network’s recurrent connections to preserve contextual information, the network predicted the next action in the sequence. Compared to the previous environments, the kitchen environment affords even greater complexity, with 21 different sequences instead of only 3. We conjectured that this more complex environment would encourage the neural networks to discover and make use of patterns that generalize across temporally extended behaviors both within and across sequences (as opposed to learning each sequence by rote). A list description of the 21 sequences is given in the appendix.

### Model architecture and training

We systematically investigated two model architectures: a non-hierarchical Elman network model, and a goal model derived from the Elman network (both models are shown in Figure 1).

**Figure 1.**
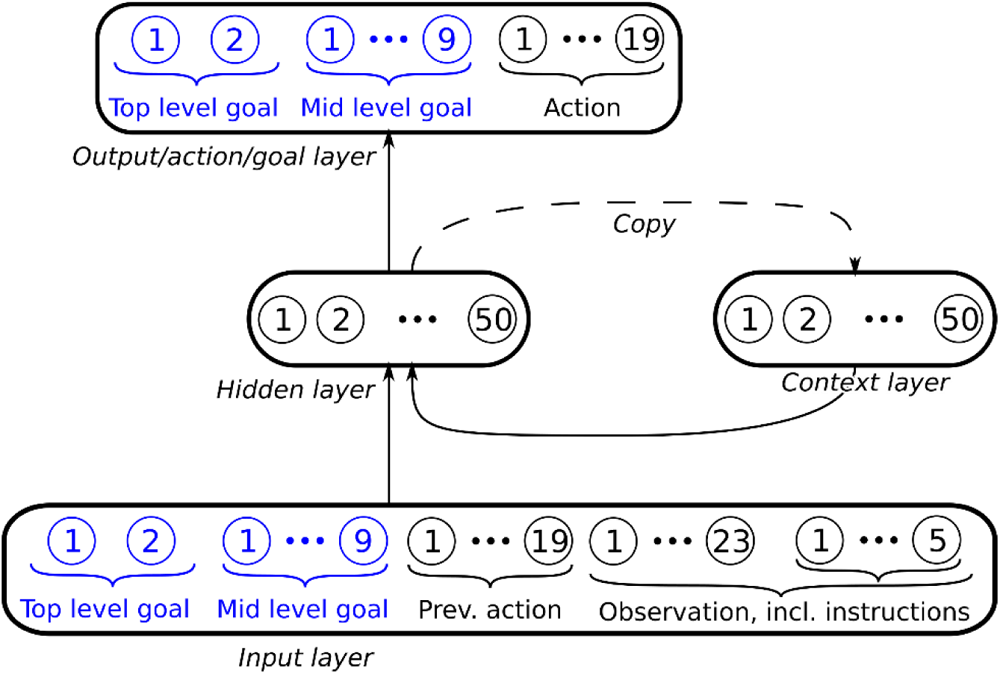
The models used for the kitchen environment task: Elman network (black) with Goal network additions (blue). See text for details.

### Model 1: the Elman Network

Elman networks are recurrent networks whereby the activation states of the hidden layer are copied to a “context” layer, the units of which serve as fully connected inputs back to the hidden layer on the next time-step (Elman, 1990). The model is shown in Figure 1 (components in black only). Its hidden layer units use the ReLU activation function. The weights were He-initialized (He, 2015) and trained using Backpropagation Through Time (BPTT) with ADAM (a popular variant of gradient descent with momentum (Kingma, 2014)), a learning rate of 0.001, and cross-entropy loss:

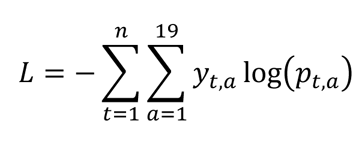

where *t* refers to the time-step out of *n* in the sequence, *a* is one of 19 possible output classes (“add grounds”, “stir”, etc.), *yt,a* equals 1 if *a* is the target action at time-step *t* and 0 otherwise, and *pt,a* is the network’s output for that action. An L2 regularization term with *k* = 10^−4^ was added to this loss. During training, sequences were sampled uniformly until maximal accuracy was achieved (with 1000 additional iterations added to ensure robustness) The networks were exposed to a sequence of inputs corresponding to a given correct behavior sequence; each step was “as if” the network had picked the correct actions at every step so far in that sequence, irrespective of what it actually chose. Furthermore, between time-steps the recurrent units corresponding to the previous action in the sequence were set to the target action for that time-step. In contrast, during the test phase the network fully interacted with the environment. This means that the actions chosen by the network were executed, which affected the unfolding trajectory of the episode; behavior deviating from the training sequences could therefore lead to previously unseen states.

To match the range of actions afforded by the environment, the network had 19 *action units* (fixate cabinet, open, etc.) in the output layer, the activity of which was, after a winner-take-all operation, included in the input layer. Other inputs consisted of 29 *observation units*, including 6 units encoding the beverage request, 15 units encoding visual observations, and 8 units encoding which object was being held. The hidden layer consisted of 50 units.

### Model 2: the Goal Network

In a second model, which was inspired by the model proposed by Cooper and colleagues (2014), we augmented the Elman network with goal and subgoal units (Figure 1, components in blue). In addition to the units of the Elman network, the Goal network included two *goal units* (representing “make coffee” and “make tea”) and nine *subgoal units* (representing “dip teabag”, “add grounds”, “add sugar”, “add milk”, “add cream”, “stir”, “tidy up”, “drink tea”, and “drink coffee”). The goal units are recurrent such that the goal unit with the maximal activation on a given time-step defines the active (one-hot) input goal on the next time-step; at any time-step *t > 0*, one goal unit, one subgoal unit, and one action unit were active. Thus, goal units were required to sustain their own activations throughout each sequence. The additional output was implemented using second and third softmax output layers for subgoals and goals, alongside the softmax layer used for actions. Thus, aside from the addition of goal units, the goal network was identical to the Elman network: it used a cross-entropy loss function and had the same number of hidden units.

During training, the (sub)goal units were set to their correct values such that they could be trained in a supervised fashion, similarly to the action units. The total loss was the sum of the cross-entropy losses for goals, subgoals, and actions:

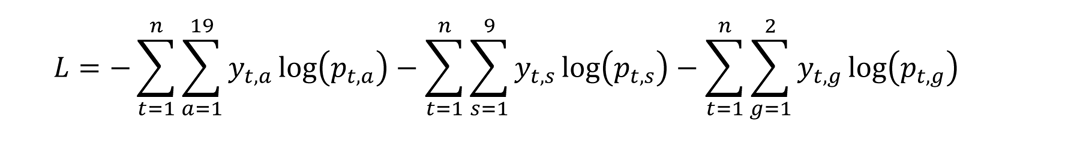

where *s* and *g* denote subgoals and goals. Training of model 2 was made as close as possible to the training of model 1, using identical initialization, training regimen, learning rate, and regularization.

### Training and task

On the first step of the task, instructions to make a specific beverage were given via dedicated input units (“instructions” in figure 1) rather than via the goal or subgoal units. For instance, the instruction to make coffee with dairy and two sugars was given by activating the units “coffee”, “dairy”, “sugar”, and “extra sugar”. These instructions were then turned off for the rest of the sequence; hence, the network was required to preserve information in its hidden layer and goal units in order to successfully create the drink.

The networks were instructed to make tea or coffee, with or without dairy, and with zero, one, or two sugars, yielding 12 possible instructions. Milk was always available and always used for tea, whereas when making coffee, cream was preferred but in some cases unavailable. As a result, there were 15 different “legal” beverages, including coffee variations with cream and with milk. Finally, the networks were trained to add ingredients in several alternative orders (sugar first or dairy first). The networks were thus trained to perform 21 different sequences in total. An example sequence is shown in table 1. Each training episode was based on one of those sequences, sampled uniformly from the 21 possible sequences.

**Table 1:** One of 21 possible sequences in the Kitchen Environment. Note that the task does not have the Markov property: ambiguous inputs force the network to preserve context across time-steps using recurrent connections in the hidden layer. This ambiguity results not only from identical observations within this sequence (e.g. step 18 and 33), but also across different sequences.

| Step | Observed | Held | Goal | Subgoal | Action |
| --- | --- | --- | --- | --- | --- |
| 1 | Make coffee, with sugar, extra sugar, dairy | Nothing | Make | Add grounds | Fixate cabinet |
| 2 | Cabinet | Nothing | Coffee |  | Open |
| 3 | Cabinet (open), coffee jar, teabags, sugar box | Nothing |  |  | Fixate coffee jar |
| 4 | Coffee jar | Nothing |  |  | Take |
| 5 | Coffee jar | Coffee jar |  |  | Open |
| 6 | Coffee jar (open) | Coffee jar |  |  | Fixate mug |
| 7 | Mug (full) | Coffee jar |  |  | Add to mug |
| 8 | Mug (full, dark) | Coffee jar |  |  | Fixate coffee jar |
| 9 | Coffee jar (open) | Coffee jar |  |  | Close |
| 10 | Coffee jar | Coffee jar |  |  | Fixate cabinet |
| 11 | Cabinet (open), teabags, sugar box | Coffee jar |  |  | Put down |
| 12 | Cabinet (open), coffee jar, teabags, sugar box | Nothing |  | Stir | Fixate spoon |
| 13 | Spoon | Nothing |  |  | Take |
| 14 | Spoon | Spoon |  |  | Fixate mug |
| 15 | Mug (full, dark) | Spoon |  |  | Stir |
| 16 | Mug (full, dark) | Spoon |  |  | Fixate table |
| 17 | Table, mug | Spoon |  |  | Put down |
| 18 | Table, mug, spoon | Nothing |  | Add cream | Fixate fridge |
| 19 | Fridge | Nothing |  |  | Open |
| 20 | Fridge (open), milk carton | Nothing |  | Add milk | Fixate milk |
| 21 | Milk carton | Milk carton |  |  | Take |
| 22 | Milk carton (open) | Milk carton |  |  | Fixate mug |
| 23 | Mug (full, dark) | Milk carton |  |  | Add to mug |
| 24 | Mug (full, dark, milky) | Milk carton |  |  | Fixate fridge |
| 25 | Fridge (open) | Milk carton |  |  | Put down |
| 26 | Fridge (open), milk carton | Nothing |  |  | Close |
| 27 | Fridge | Nothing |  | Stir | Fixate spoon |
| 28 | Spoon | Nothing |  |  | Take |
| 29 | Spoon | Spoon |  |  | Fixate mug |
| 30 | Mug (full, dark, milky) | Spoon |  |  | Stir |
| 31 | Mug (full, dark, milky) | Spoon |  |  | Fixate table |
| 32 | Table, mug | Spoon |  |  | Put down |
| 33 | Table, mug, spoon | Nothing |  | Add sugar | Fixate sugar box |
| 34 | Sugar box (open) | Nothing |  |  | Take |
| 35 | Sugar box (open) | Sugar cube |  |  | Fixate mug |
| 36 | Mug (full, dark, milky) | Sugar cube |  |  | Add to mug |
| 37 | Mug (full, dark, milky) | Nothing |  |  | Fixate sugar box |
| 38 | Sugar box (open) | Nothing |  |  | Take |
| 39 | Sugar box (open) | Sugar cube |  |  | Fixate mug |
| 40 | Mug (full, dark, milky) | Sugar cube |  |  | Add to mug |
| 41 | Mug (full, dark, milky) | Nothing |  | Tidy up | Fixate cabinet |
| 42 | Cabinet (open), coffee jar, teabags, sugar box | Nothing |  |  | Close |
| 43 | Cabinet | Nothing |  | Stir | Fixate spoon |
| 44 | Spoon | Nothing |  |  | Take |
| 45 | Spoon | Spoon |  |  | Fixate mug |
| 46 | Mug (full, dark, milky) | Spoon |  |  | Stir |
| 47 | Mug (full, dark, milky) | Spoon |  |  | Fixate table |
| 48 | Table, mug | Spoon |  |  | Put down |
| 49 | Table, mug, spoon | Nothing |  | Drink coffee | Fixate mug |
| 50 | Mug (full, dark, milky) | Nothing |  |  | Take |
| 51 | Mug (full, dark, milky) | Mug |  |  | Sip |
| 52 | Mug | Mug |  |  | Fixate table |
| 53 | Table, spoon | Mug |  |  | Put down |
| 54 | Table, mug, spoon | Nothing |  |  | Say 'Nice coffee!' |

Because of random weight initialization and random ordering of the training samples, neural networks frequently show substantial individual variations even when trained using the same hyperparameters. Accordingly, 50 networks were trained for each architecture to ensure that the results were robust; training time varied between 4,000 and 16,000 iterations.

### Experiments on the model

Hierarchy allows for modular and compositional manipulation of low-level behaviors and may help produce representations of abstract task goals that are robust to variations of superficial features of the task. We investigated whether the goal units enhanced or degraded the goal network’s performance when stressed in novel conditions. First, in order to evaluate and compare the contribution of the different model components, we conducted ablation studies^1^ by deactivating some categories of units. Second, to examine whether hierarchical structure resulted in unique patterns of behavior when stressed, we added Gaussian noise to the hidden unit activations. Third, to test whether goal-unit activity could elicit classic signatures of cognitive control from the behavior of the model, we modulated the activation states of the goal units in search of evidence for top-down suppression of habitual actions or for facilitation of non-habitual actions.

We use the following metrics throughout the following sections:

- Errors are defined as the first deviation from correct behavior on each trial (as subsequent deviations on the same trial can be caused by the network entering never-seen-before states of the environment resulting from the first deviation).
- We distinguish two types of errors: “subsequence errors” and “action errors”. Subsequence errors are errors whereby one subsequence is replaced by another subsequence, for instance adding cream instead of adding milk, or skipping “add sugar” and instead drinking the unsweetened beverage. In some cases, a subsequence error leads to the creation of one of the possible beverages but not the one requested. Action errors are any errors that are not subsequence errors; for example using the action “stir” instead of “open” when fixating the milk carton.

#### Ablations

In this first series of interventions, we simulated damage to the network by successively turning off the goal, subgoal, action, and observation units in order to investigate the degree to which the network relied on each of those unit types for its performance.

Figure 2 illustrates the results. Notably, even when the observation units were inactivated (rendering the network effectively blind) the goal network completed at least one subsequence in 18% of sequences, based solely on the neural dynamics of the recurrent units (Figure 2, “observations”, orange bar). With other ablations, the goal network was able to complete some complete sequences (Figure 2, beige bars). Ablating the action units, corresponding to immediate one-step temporal relationships, induced a greater proportion of action errors (Figure 2, “actions”, blue bar), whereas ablating the goal and subgoal units induced a greater proportion of subsequence errors in erroneous sequences (Figure 2, “Subgoals”, “Goals”, and “Subgoals and Goals”, orange bars). This suggests that, in the goal network, goal units played an important role in representing temporally extended, abstract patterns of behavior.

**Figure 2.**
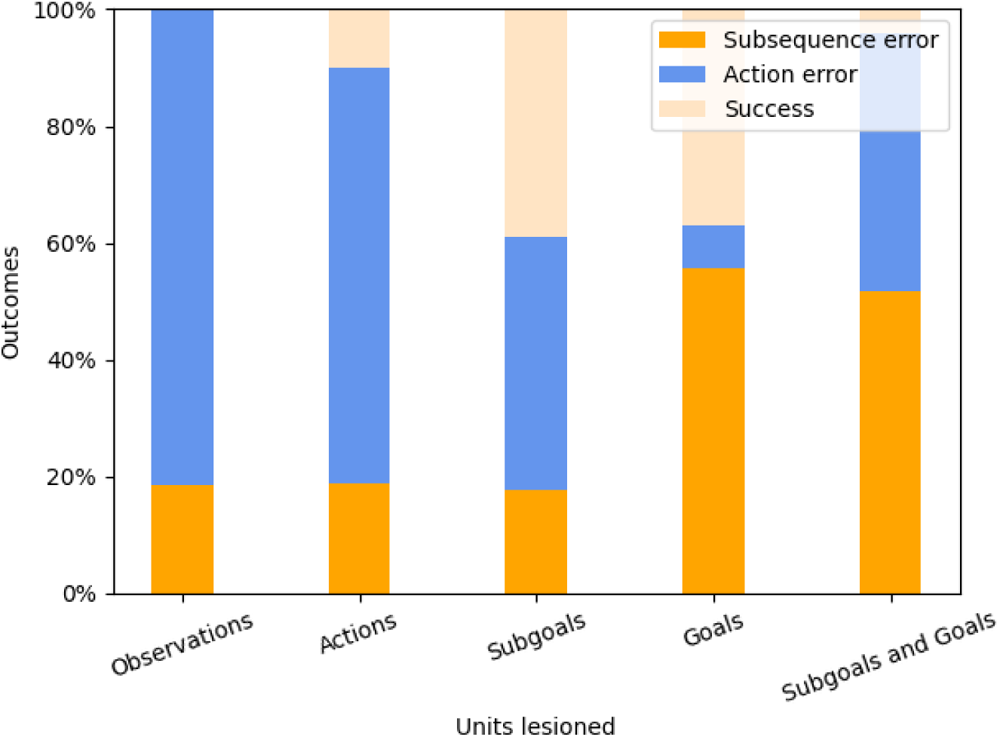
Ablation studies: performance of the network when different groups of units are lesioned (all connections set to 0). Outcomes are separated into subsequence errors (the first error encountered consists in substituting one subsequence with another, e.g. adding sugar instead milk), action errors (any other error), and success (no errors). These manipulations show that the network makes use of all the units to perform the task, and confirms that lesioning more abstract units leads to more structured errors behaviorally.

#### Stressing the model with noise

In the second category of interventions, we simulated dynamic disturbances that could affect task execution: distractions due to unrelated tasks and perceptual uncertainty. Following Botvinick and Plaut (2004), distraction was simulated by adding Gaussian noise to the hidden units of the model (before these values were put through the activation function). To study the effects of noise in a systematic manner, each network was run 10 times on each of the 21 sequences. This was repeated for each noise magnitude.

##### Qualitative assessment

We expected that most errors would occur at the inflexion points of the behavior, i.e., on steps on which the network begins to work towards a different subgoal (e.g., adding a sugar, stirring, etc.), as previously observed by Botvinick and Plaut (2004). This behavior would be consistent with the network implicitly capturing the task structure. This was indeed the case, as illustrated by Figure 3 for the sequence “coffee with cream and sugar, sugar first”. As can be seen by inspection, the errors mostly occurred on the first few actions of a sequence and at the transition points between subsequences. Nevertheless, it is noteworthy that the pattern of errors did not exactly correspond to the task subsequences; for instance, the more frequently occurring and perhaps more predictable “stir” sequences elicited fewer errors.

**Figure 3.**
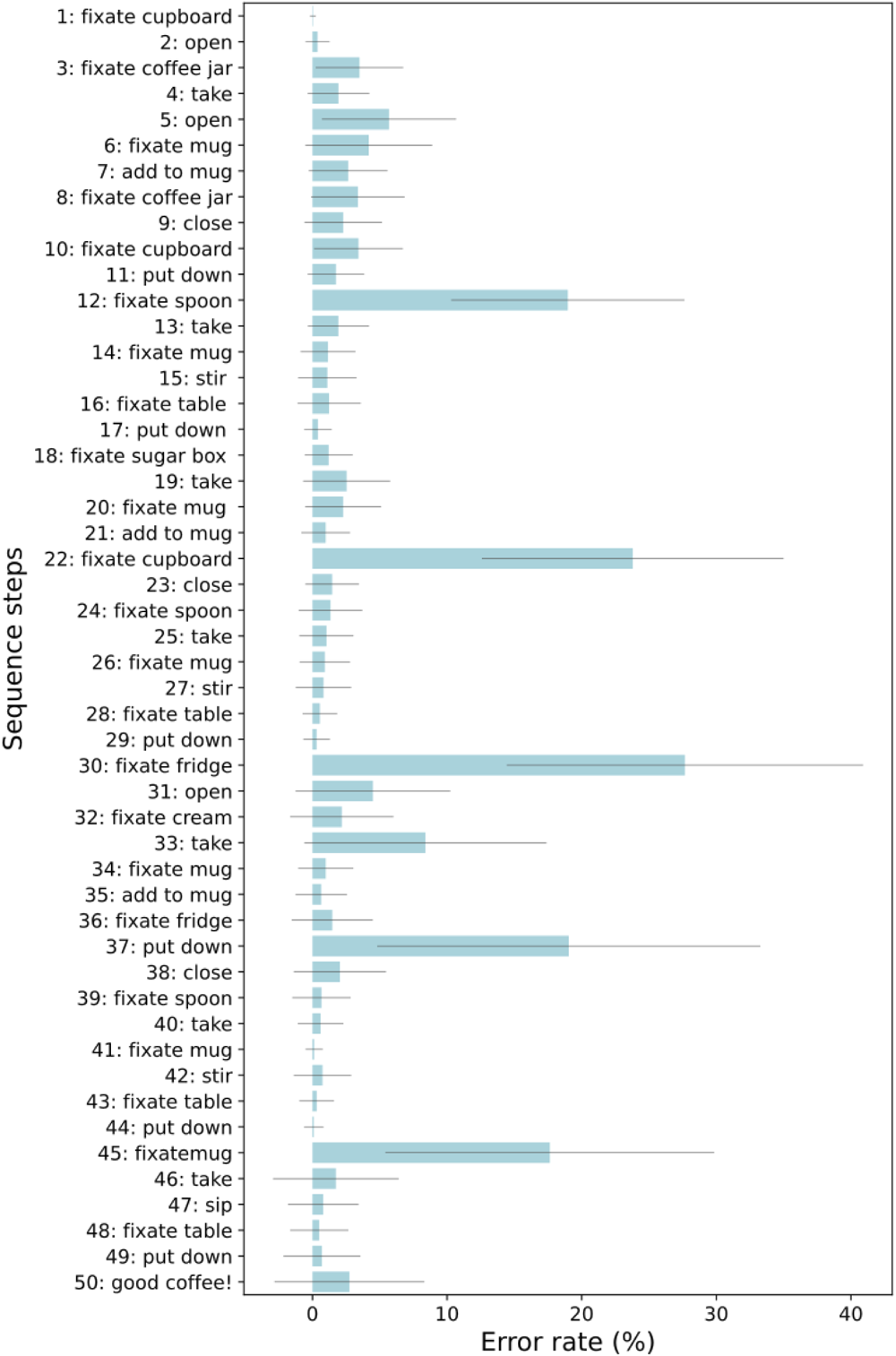
Network error rate under stress, with standard deviations in black. All 50 networks were tested for a 100 runs each on the 50-step sequence “Coffee with sugar and dairy”. The vertical axis indicates each step of the sequence. Gaussian noise (mean 0, magnitude 0.25) was added to the pre-activation values of the hidden layer at every step. Note that steps corresponding to transitions between sub-sequences are often more error-prone, e.g., step 12 begins a “stir” subsequence, step 22 begins a “tidy-up” subsequence (in which the cupboard is closed as no more ingredients from inside it are needed), step 30 begins an “add cream” subsequence, and step 45 begins the “drink coffee” subsequence. However, this was not always the case: stir subsequences (steps 12-17,24-29 and 39-44) had a lower error rate overall, presumably because they were the most common subsequence type and almost always followed immediately after an ingredient was added, making them easier to carry out.

##### Quantitative assessment

To study the effects of this intervention systematically, we counted the total number of errors and the proportion of subsequence errors versus action errors, separately for the Elman network model and the goal network model. We expected that task structure incorporated by the hierarchical model would limit the degree to which the noise degraded its performance relative to the performance of the Elman network. In particular, we expected the goal networks to produce a greater proportion of subsequence errors as compared to the Elman networks. By more strongly distinguishing between subsequences, the goal representations were expected to restrain switches away from the currently active subsequence until the end of that subsequence. Instead, representations of *future* subsequences might be more susceptible to noise, causing the errors to occur later at the transition points between the subsequences (see, e.g., Botvinick & Plaut, 2004; Botvinick & Bylsma, 2005).

Figure 4 illustrates the networks’ behavior in the presence of varying levels of noise added to the activation states of the hidden layer units. Predictably, for both networks (solid and hashed bars), increasingly higher levels of noise resulted in increasingly worse performance (1-5, pink bars), indicating that the noise generally destabilized the networks. Further, with low levels of noise a higher proportion of these errors were subsequence errors (0.01 to 0.5, red bars), whereas with higher levels of noise the errors were mostly action errors (1.0, orange bars). However, although incorporation of the goal and subgoal units improved the robustness of the network against noise somewhat, the difference in the proportion of subsequence vs. action errors between the action and goal models was small (less than 10% at intermediate noise levels; compare the solid vs. the hashed bars). These results suggest that, contrary to our expectations, the incorporation of goal units did little to improve the performance of the network under stress, nor its ability to capture temporal structure in the task.

**Figure 4.**
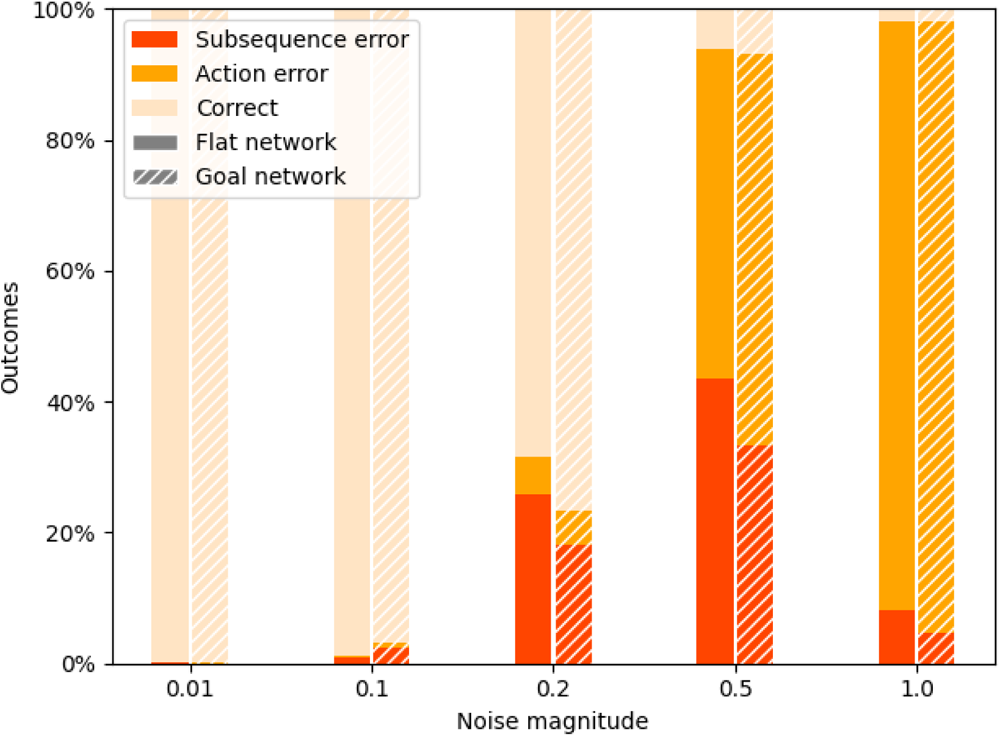
Relative percentage of errors types in an example sequence as a function of noise magnitude, for the flat network and goal network. The x-axis indicates the level of Gaussian noise added to the input to the hidden layer units (prior to the application the non-linearity). Noise magnitude is given in arbitrary units.

To investigate how noise affected the network’s function over time, we also added noise to the hidden unit activations for a single time-step on each trial of each simulation. This enabled us to measure the length of the interval between the external perturbation and its consequences on the network’s overt behavior. Specifically, each sequence was run as many times as there were steps in that sequence, such that the effect of adding noise on each separate time-step could be observed in isolation. This was repeated for each of the 50 Elman networks and each of the 50 goal networks, at Gaussian noise magnitudes of 1.0, 2.0, 3.0, 4.0, and 5.0; we recorded the number of time-steps between when the noise was added and the first deviation in the sequence, separately for action errors vs. subsequence errors. Figure 5 displays the results for a noise magnitude of 1.0: the action errors consistently followed immediately after each perturbation, whereas the subsequence errors tended to occur after longer delays, for both the Elman and Goal networks. At higher noise magnitudes, the effects remained but the delays tended to be shorter, for both network types (e.g., for the Goal network at magnitude 5.0, the average delays were 0.39 and 1.37 for action and subsequence errors respectively; full results are reported in the appendix).

**Figure 5.**
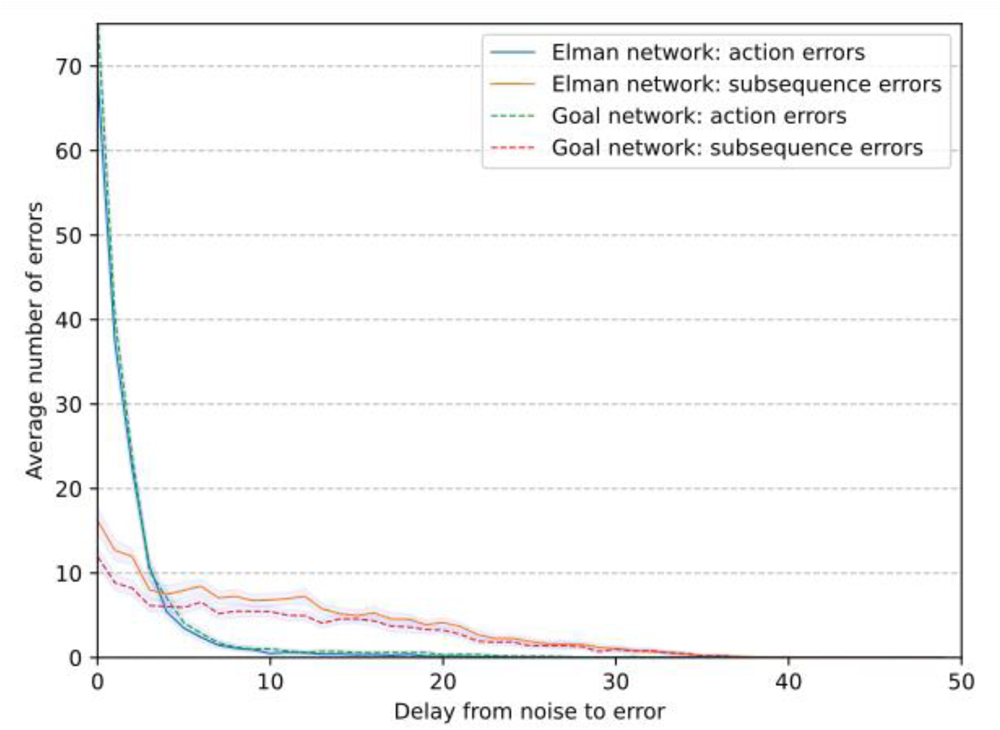
Delay (number of steps from noise perturbation to error commission) for Gaussian noise at magnitude 1.0.. We compare the Goal and Elman networks separately for subsequence errors and actions errors. Following the introduction of noise, subsequence errors tend to occur with a longer delay relative to action errors, which occur immediately after the noise. 95% confidence intervals are shown in grey.

#### Control

In a third series of interventions, we investigated whether the goal units could steer the behavior of the goal network towards desired end states. Specific goal units in the goal model were activated or deactivated in order to facilitate or inhibit the production of the corresponding action sequences. Furthermore, to override the network’s trained behavior, we multiplied the goal unit activations by a factor >1 (see Table 2), thereby applying a level of control sufficient to alter behavior.

**Table 2:**
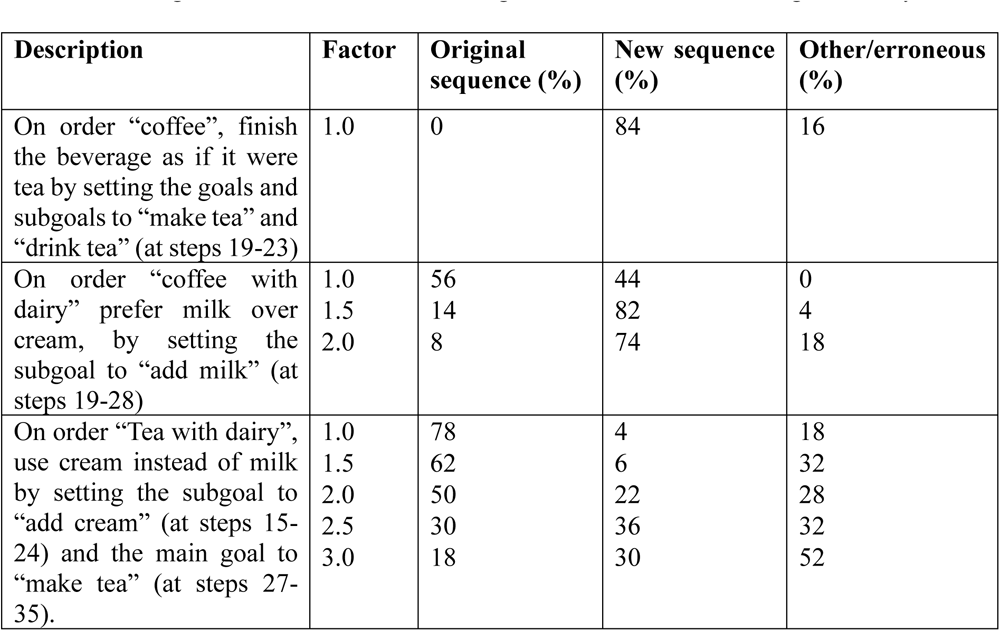
Using the goal units to control behavior. All 50 networks are tested in each condition. We report the proportion of networks whose behavior remains unaffected (original sequence), that produces the expected sequence by behaving according to the activated goals (new sequence), and that produces some other sequence (other/erroneous). For each condition, the goal units were activated at the same values used during training, or were multiplied by a scalar (Factor) to increase the effect of the units on the network’s behavior. The results show doing so increases the probability of producing the intended new sequence but also causes more errors. Such top-down modulation can overcome a well-trained preference (e.g., during training, the network always used cream with coffee, when cream was available) and to produce entirely new sequences by recombining subsequences (e.g., making tea with cream, despite the network never being trained on this sequence).

##### Steering the network

Cognitive control enables agents to overcome habitual actions in order to perform rare or novel behaviors. Here we asked whether application of top-down control could guide the production of novel actions sequences. First, we tested the models on a simple modification of the task affecting only its last action: saying “good tea” instead of “good coffee” at the end of a “make coffee” sequence. To do so, we activated the “make tea” goal and “drink tea” goal at the end of the sequence. Second, we investigated whether the goal model could be induced to add milk instead of cream to coffee, even when cream was present, despite the model being trained to add milk only when cream was not available. Accordingly, for the coffee-making sequences where cream was present, we activated the “add milk” subgoal unit instead of the “add cream” subgoal unit of the goal model, and further manipulated the strength of this effect by increasing the goal unit activations by a constant factor. Finally, we used the same technique to steer the network into producing a completely novel beverage sequence, tea with cream.

These results are shown in Table 2. Although these interventions sometimes produced errors, they were often capable of producing original and valuable sequences without any additional training. In particular, although we were concerned that recurrent connections could lead to instability, this was rarely the case. The strength of the intervention (column 2 in Table 2) could be used to manage the trade-off between forcing new behavior and avoiding instability.

We used t-SNE (Van der Maaten & Hinton, 2008) to visualize the effects of changing goals on the hidden layer representations when making tea with cream instead of with milk. Three sequences are shown: the training sequences tea with milk and coffee with cream, and the original sequence tea with cream (Figure 6). Note that all three sequences begin at the center of the figure and end at the bottom left. The yellow (“make coffee with cream”) and red (“make tea with milk”) trajectories correspond to the execution of trained sequences, whereas the blue (“make tea with cream”) trajectory corresponds to the execution of the novel tea-making sequence. Note that the patterns of activation for the “tea with cream” sequence are very similar to those of the “tea with milk” sequence at the start, including dipping the tea bag and stirring. By contrast, the “tea with cream” trajectory departs from the “tea with milk” trajectory and approaches the “coffee with cream” trajectory upon activation of the “add cream” subgoal. In so doing, the network generates a new sequence by recombining the subsequences from two different beverage recipes.

**Figure 6.**
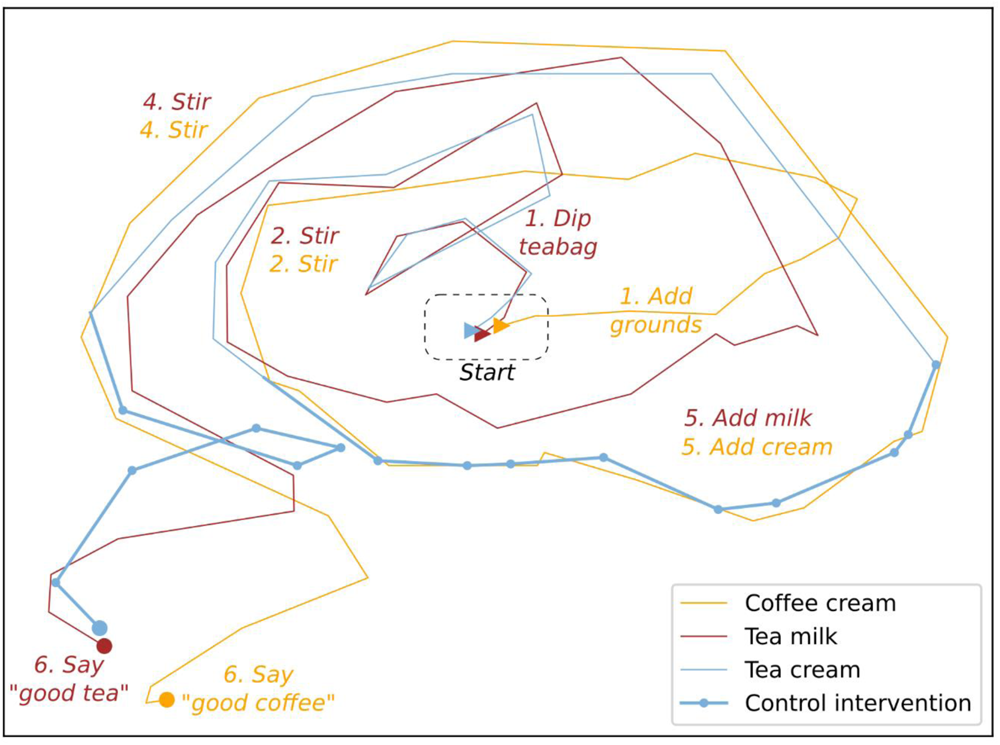
the effect of control on hidden layer representations, visualized using t-SNE. Two sequences (Coffee cream and Tea milk) are part of the training set, whereas the third sequence, Tea cream, was not in the training set and was obtained instead by manually setting goals and subgoals to drive the network (“Control intervention” steps, corresponding to the last row of table 2). The Tea cream sequence received the same instructions as the Tea milk sequence (“tea with dairy”) and followed an identical representational trajectory, until the activation of the “add cream” goal drove the network state closer to the representations associated with “coffee cream”. Subsequently the network followed a similar trajectory to “coffee cream” until activation of the “make tea” goal ensured a correct completion of the sequence.

##### Control and representations

To understand better how goals units shape network behavior and the relationship between hierarchy and distributed representations, we measured the representational changes occurring in the hidden layer when goal units activations were multiplied by a constant. As illustrated above, a low-dimensionality visualization (Figure 6) provides some intuition as to the forcing effect of goal units on hidden layer representations: over multiple time-steps, the goal units moved the representational trajectory towards the representational space associated with those goals.

Toward this end, we varied the magnitude of goal unit activations, multiplying them by 0 (i.e., de-activating those units), 1 (similar to the training regime), or 2 (doubling their activation). Figure 7 uses t-SNE to illustrate the effect of this intervention for the first 6 steps of two sequences (Coffee without dairy or sugar, and tea without dairy or sugar). There were relatively few errors, at least on these first few actions, even when goal units were fully deactivated. Indeed the distributed representation of the task enabled the network to pick the correct behavioral trajectory even without control; greater goal unit activations caused only slight deviations from this representational trajectory on these short sequences (though errors did eventually occur over the course of the full sequence; see figure 1). Secondly, the goal units appear to play a role in “shifting” the tea and coffee representations apart, such that deactivating the goal units results in more similar representations for the tea versus coffee sequences, whereas doubling the activation of the goal units results in representations that are further apart, especially near the beginning of each sequence. For instance, for step 2 of each sequence in Figure 5, the distance is maximal between “coffee x2” and “tea x2” and minimal between “coffee x0” and “tea x0”.

**Figure 7.**
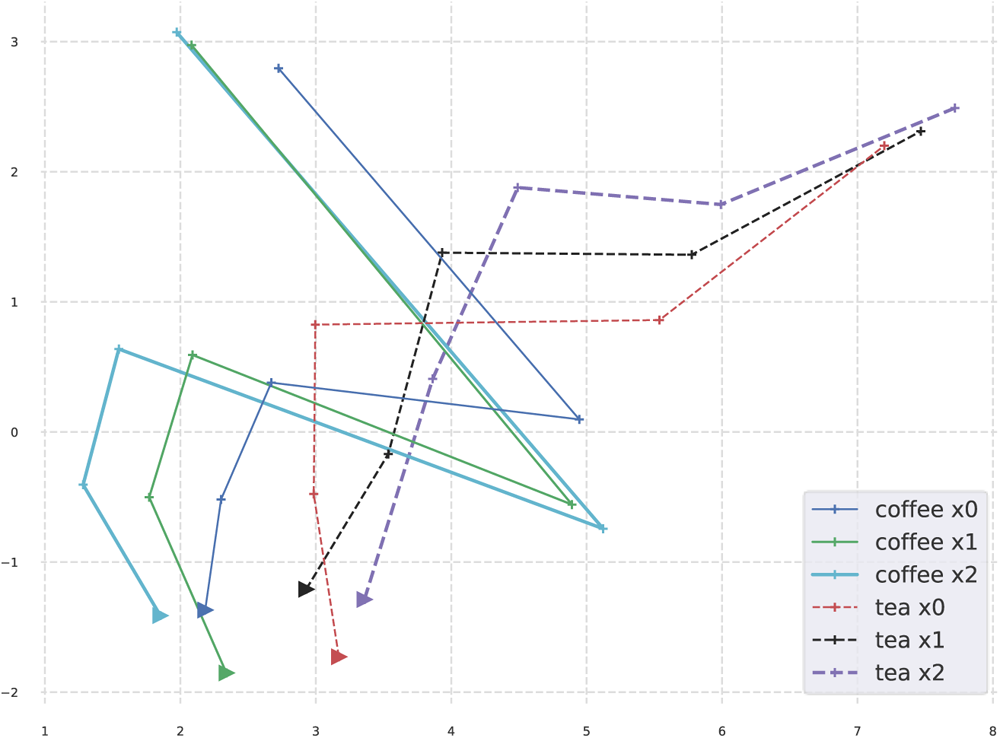
t-SNE visualization of the activations of the hidden units for the first five steps of 6 sequences: making tea or making coffee, with goal multipliers set to 0, 1, or 2. At x0, the goal units are deactivated; at x1 they behave as in their training; and at x2, their activation is doubled. The sequence origins are indicated by a triangle. The x0 sequences are close to one another whereas the x2 sequences are furthest apart, showcasing how goals increase representational dissimilarity between actions corresponding to different goals and subgoals. Note that this effect is especially strong on the initial actions of a sequence.

A systematic measurement of the Euclidian distances between hidden layer activations for all 50 networks, over all time-steps of all sequences, confirmed that goal units contributed to increasing the distances between the representations of actions aiming to accomplish different goals. Multiplying goal activations by 2 increased representational distance by 0.033 for activations that shared the same goals and subgoals, compared to 0.061 when either the goal or the subgoal was different, and 0.091 when both the goal and subgoal were different (all comparisons significant, p<0.001, t-test). This confirms the qualitative shift observed in Figure 7, though it should be noted that the effect is small relative to overall representational distances (as the average Euclidian distance between any two representations, without any intervention, is 14.1), and is likely magnified in t-SNE visualizations (which emphasize the preservation of local structure over long-range order).

### Study 1 summary

Comparison of the Goal and Elman networks revealed that the latter displayed considerable functionality in the execution of extended action sequences despite the lack of hierarchical representations. The Elman networks were nearly as robust to noise relative as the hierarchical networks (Fig 4). Not only was the accuracy of the Elman network comparable to the Goal network under stress, but their behavioral degradation was characterized by similar structure: in both cases, errors tended to consist of extended subsequences matching other task components – so called “action slips” – rather than random actions. This supports previous claims that recurrent networks can capture relevant task structure needed to carry out extended sequences, even without incorporating hierarchical representations (Botvinick & Plaut, 2004).

Nevertheless, rather than improving the robustness of the network when performing familiar tasks under stress, we observed that the goal units could flexibly regulate network behavior for novel tasks. Indeed, the goal network could sometimes succeed even on the first trial of a new task, namely by exploiting new combinations of learned behavior patterns as guided by corresponding subgoals. Higher activation of the subgoals was more likely to modify the previously-learned (reflex) behavior, albeit with a higher probability of making an error.s We then compared representations in the goal network to those in the Elman network. Additionally, within the goal network, we investigated the difference between the standard representations and the representations when the goal unit activations were perturbed, both by inactivating them and by multiplying them by a constant factor. These results are elaborated in the general discussion.

### Study 2: Comparing the hierarchical model with fMRI data

The preceding simulations suggest one mechanism whereby recurrent neural networks can balance the costs and benefits of hierarchy vs. distributed representations. Here, we investigated how the brain regulates this trade-off, especially regarding frontal brain areas, which are thought to implement cognitive control according to spatially-organized hierarchical representations (Badre & Desrochers, 2019, Christoff et al., 2009). We asked whether and how the goal model might relate to brain areas associated with cognitive control. Toward this end, we re-trained the Elman and Goal models on a “coffee-tea making” task, which is a hierarchical sequence task that was previously used with human participants in an fMRI experiment (see Holroyd et al., 2018). In addition, we tested a third model that was trained with a loss function that penalizes long-distances connections (“wiring costs”, as described in the following). After training, representational similarity analyses (RSA) were used to compare the representations of these three models during the sequence production task against those of the previously collected human fMRI data set (Holroyd et al., 2018), allowing for direct comparisons between the different models.

### The coffee-tea task

Participants were instructed to assume the role of a barista preparing tea and coffee for their customers. Notably, they themselves choose which beverage to prepare, but were instructed to do so randomly as if “flipping a coin”. The preparation of each beverage always took six action steps. During each step they saw three images (e.g., during step 1: coffee beans, tea grounds, and pepper) from which they were required to select one by pressing a spatially-corresponding button on a button box (Figure 8). If they prepared a beverage correctly or incorrectly they were shown a happy or sad smiley as visual feedback, respectively, representing a happy or sad customer.

**Figure 8.**
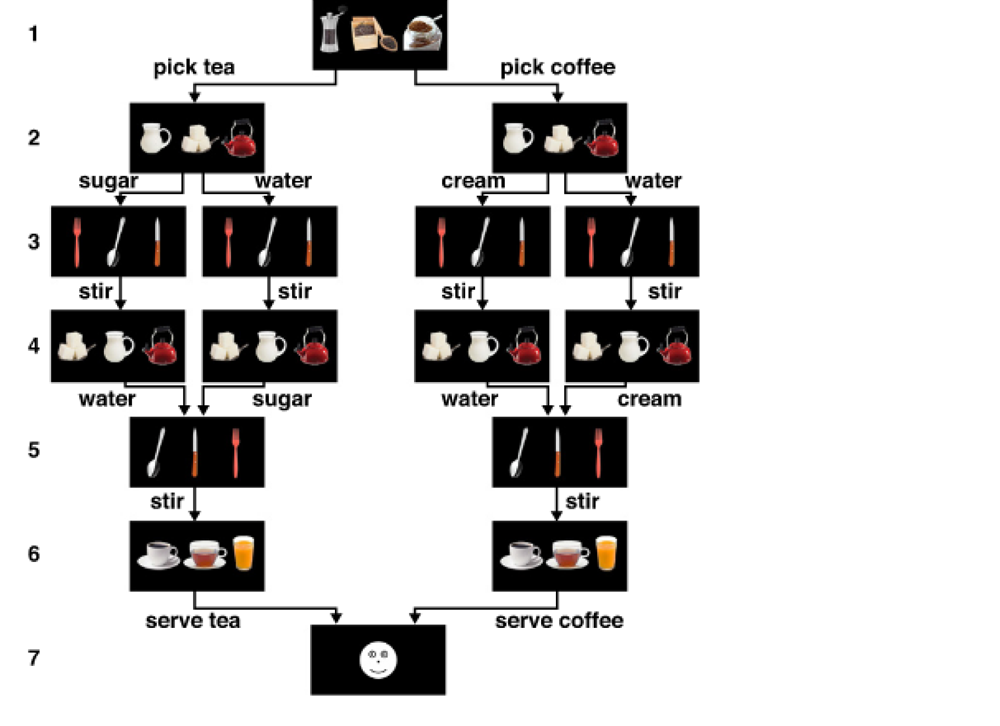
the coffee-tea task

**Figure 9.**
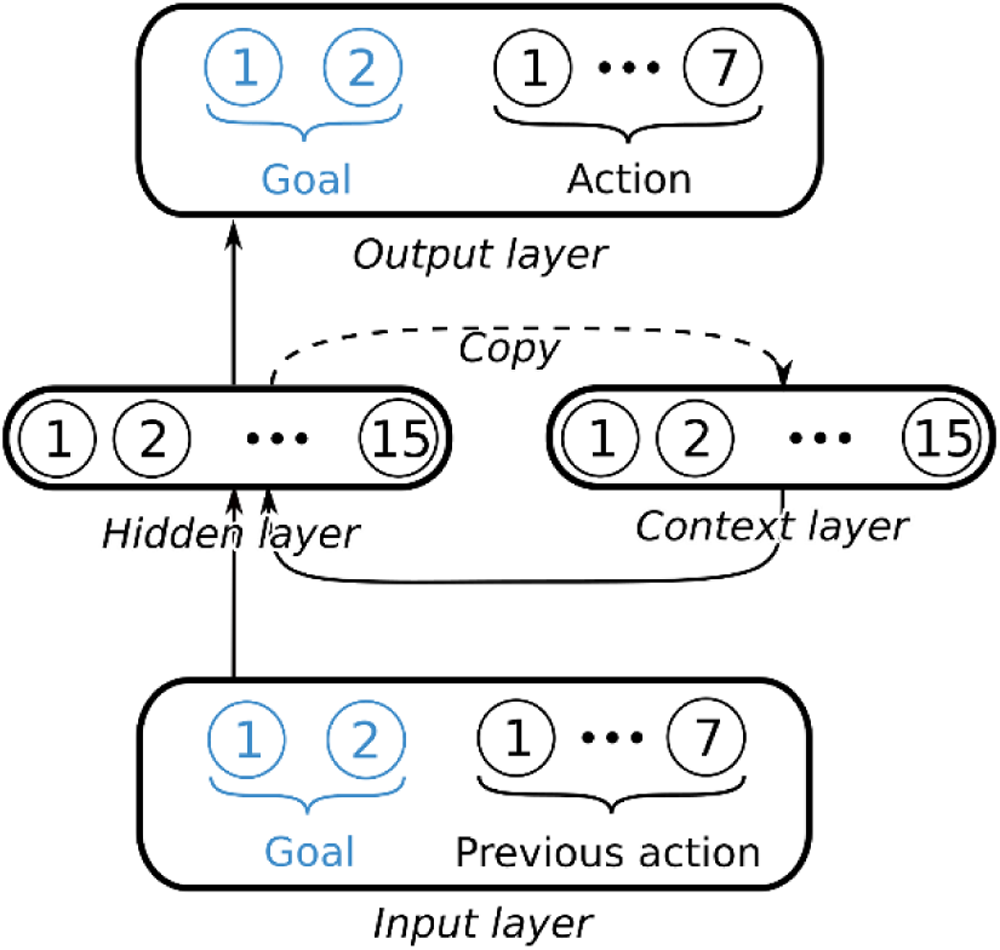
Neural networks used for the coffee-tea task. In black, the Elman network; in blue, goal units added in the Goal network.

The task rules resulted in 4 possible correct task sequences: two for preparing coffee (by either adding water first and then cream, or adding cream first and then water) and two for preparing tea (by either adding water first and then sugar, or adding sugar first and then water). In figures these sequences are abbreviated as “cw1” (coffee, water first), “cw2”, “tw1”, and “tw2” respectively. Participants were instructed to decide at random which ingredient (water or (depending on the beverage) cream or sugar) to add first. Because participants saw the same images on action steps requiring different responses (although the position of each image varied both within and across action steps), they needed to remember the context in which the actions were selected. In particular, they were required to remember what they were preparing (coffee or tea), and which ingredient they had added first (cream, sugar, or water).

With 4 correct task sequences that each consisted of 6 action steps, the task included in total 24 unique action steps. The three images displayed during each action step were shown to participants for 1s, followed by a black screen for 3.5 +/-1s (with 7 possible jitters at .33s increments). If no response was given before the three images of the next step appeared, the words “too slow” appeared and the trial was rendered incorrect. Feedback (happy/sad smiley) was also shown for 1s, followed by a 3.5 +/-1s ITI. Full details of the task are given by Holroyd et al. (2018).

### Neural Network Modeling

We extended our original RNN model of sequential task execution in a series of steps that gradually increased the degree of hierarchy incorporated in the model, in order to examine the progressive impact of these changes on ACC representations. The resulting 3 networks yield hidden layer activation data that can be compared with multivariate brain BOLD data from fMRI using RSA. Toward this end, we simulated a version of the coffee-tea task used previously in an fMRI experiment (Holroyd et al., 2018; see also Fig. 8), as shown for one sequence in Table 3. Each of the 3 RNNS was then trained to execute this task, while the fMRI participants did this task while lying in the scanner.

**Table 3:** One of four possible sequences in the coffee-tee making task.

| Step | Input | Goal | Action |
| --- | --- | --- | --- |
| 1 | Start | Make coffee | Add grounds |
| 2 | Add grounds |  | Add water |
| 3 | Add water |  | Stir |
| 4 | Stir |  | Add sugar |
| 5 | Add sugar |  | Stir |
| 6 | Stir |  | Serve coffee |

We then constructed per RNN a representational dissimilarity matrix (RDM) that indicates how (dis)similar the activity patterns of the hidden layer were across all pairwise combinations of task states for, in this case, the 24 unique steps of the coffee-tea task (Kriegeskorte, Mur, & Bandettini, 2008). For instance, the activity patterns across the hidden layer during a stirring action may be more similar to the activity patterns for a different stirring action than to those corresponding to an add-water action. Likewise, using a whole-brain searchlight approach, we constructed brain RDMs that indicate how (dis)similar the activity patterns were across a group of voxels across all 24 pairwise combinations of task states. Crucially, creating RDMs provided a means for comparing second-order similarities between these representations encoded in disparate sources of data (e.g., hidden unit activation values in an RNN vs. voxels of fMRI data). Model RDMs that were strongly correlated with brain RDMs indicate a strong correspondence of their representations across conditions: the representations reveal second-order similarity.

As a basis for comparison with the other two networks, we first replicated the results of the original RNN model used by Holroyd et al. (2018), which was an Elman network incorporating recurrent connections that maintained the task context as the sequences unfolded.

Next, we added goal units to the model that explicitly distinguished between the two tea and two coffee sequences. At the start of each trial, the goal units were set to “tea” or “coffee” according to the type of target sequence. This goal-related information was then maintained by the distributed pattern of activations in the hidden layer, supported by the recurrent connections as the trial progressed. In line with a theory of medial frontal cortex (Holroyd & Yeung, 2012; Holroyd & McClure, 2015; Holroyd & Verguts, 2021), we predicted that application of RSA to the representations in the hidden layer of the goal network would resemble activity patterns in rostral areas of ACC, corresponding to more abstract task representations.

Third, by incorporating an additional cost into the loss function during training, we simulated wiring costs that induced an abstraction gradient in the hidden layer (see below). This enabled us to map different parts of the model’s hidden layer to different regions of the ACC, and ask whether ACC exhibits a similar spatial gradient.

#### Model 1 and 2: Adaptations of Elman and Goal networks for the fMRI task

As described above, the RNNs used in the kitchen environment in Study 1 were adapted to the relatively simple task developed for use in fMRI experiments. The hidden layer had 15 units (down from 50), with a sigmoid activation function. We used cross-entropy loss, a learning rate of 0.1, and no L2 regularization. The networks were trained by stochastic gradient descent, using backpropagation through time, over the 4 possible sequences. Note that, for the Elman network, the only departure from the training procedure utilized by Holroyd et al. (2018) was the use of a cross-entropy loss, instead of mean square error. All of the networks were trained for 5000 sequences, which was sufficient to reach perfect (100%) accuracy. To ensure replicability of the results, for each model we used the mean RDM from 100 networks.

The difference between the Elman and Goal networks consisted in the introduction of two goal units, corresponding to the goals “tea” and “coffee”. Similar to the models used in study 1, we used separate softmax layers for the different outputs (goals and actions); in contrast with those models, there were no subgoals in this experiment.

At the beginning of each sequence, the recurrent layer activations were set to 0; then as the sequence progressed the input states (“Start”, etc.) were given to the network as one-hot vectors.^2^ Note that, following Holroyd et al. (2018), the input at each time-step corresponds to the previous output action (except for “Start”), simulating short-term memory rather than the stimulus itself.

#### Model 3: the abstraction gradient goal network

Our analysis so far has focused on the computational tension between hierarchical vs. distributed representations. However, in real brains, these computational differences give rise to an actual physical tension: neurons occupy space and are physically distant from one another, which results in significant energetic costs that depend mainly on the length of axons between distal brain regions (Bullmore & Sporns, 2012, Chklovskii & Koulakov, 2004, Chen et al., 2006). Accordingly, we hypothesized that hierarchically-high mechanisms involved in goal regulation and low-level systems closer to action production might be subjected to similar biophysical constraints. To explore this possibility, we created a third model in which this trade-off was enforced by way of the loss function. As before, we assumed that the hidden layer spanned the distance between the action units and the goal units, representing ACC. We further assumed that the model units were spatially organized, such that action units (corresponding to caudal and dorsal areas of ACC close to the supplementary motor area) were located towards one side of the hidden layer, and the goal units (corresponding to more rostral areas of ACC) were located on the other side. To implement this spatial organization, a loss term was added to the loss function as follows. If wiring costs are proportional to the distance between units and the strength (weight) of the synaptic connections, then the additional loss term can be expressed by the following equation, where *wi,j* denotes the weight of the recurrent connection between the *i*th and *j*th units of the hidden layer (out of a total of *n* units – n=15 in this study):

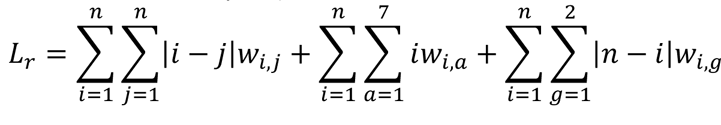

The first term corresponds to the loss incurred by recurrent connections (which penalizes communications between distant neurons). The second term corresponds to the loss incurred by connections between the hidden layer and action output units, assuming that unit 1 is closest to the action units of the output layer, and unit *n* is furthest from them. Finally, the third term corresponds to the loss incurred by connections between the hidden layer and goal output units, assuming that unit *n* is closest to the goal layer and unit 1 is furthest from it. This estimation of wiring costs is consistent with the wiring costs literature (Cherniak 1994, Chenet al. 2006).

Owing to this manipulation, units 1 to *n/2* of the hidden and context layer were incentivized to have stronger connections to goals, and units *n/2+1* to *<u>n</u>* were incentivized to have stronger connections to actions, inducing a gradient of abstraction along the hidden layer. In order to conduct the RSA, we generated separate RDMs for the ‘goal’ hidden layer units (the 7 units closest in wiring distance to the goal output units) and the ‘action’ hidden units (the 8 units closest in wiring distance to the action output units). From these we produced two different RDMs: an ‘action’ RDM and a ‘goal’ RDM. As with models 1 and 2, the final RDMs were created by averaging the distances across the respective 100 RDMs associated with each of the trained networks, separately for each element of the RDM.

#### Hyperparameter robustness exploration

The process of neural network design involved choices among many hyperparameters that could affect the performance and behavior of the network. For the sake of simplicity and to minimize the degrees of freedom associated with hyperparameter tuning, our models used the hyperparameters in Holroyd et al. (2018) (see appendix II in that article). However, we were concerned about whether the results would remain robust against variations in the hyperparameters. To mitigate this concern we trained multiple Elman networks with different sets of hyperparameters. In particular, we varied L1 and L2 regularization (k=0.0001 or k=0 for each), the activation function (ReLU, sigmoid, or tanh), the number of hidden units (8, 15, or 50), the loss function (cross-entropy or mean square error) and the weight initialization (using uniform or normal distributions). The learning rate was used as an adjustment variable to ensure that at least half of the networks for each network architecture achieved maximal accuracy in all conditions. 100 networks, with varying initialization values, were trained for each of 144 different hyperparameter combinations (14400 networks in total). An average RDM was produced from all those networks and a searchlight RSA was conducted using that RDM.

Note that because of an exhaustive search over the parameters, many of these networks occupied remote regions of the hyperparameter space associated with suboptimal task performance (e.g., for the networks that combined inefficient initialization and activation functions with a low number of units and double regularization). The resulting average RDM was therefore intended as a sanity check: we expected that it would be less sensitive to brain representations than the RDMs produced by the Elman network described above, but if these neural network representations are robust to variations in the hyperparameters, then RSA ought to select similar areas for this average RDM as for the Elman network.

### fMRI methods

We re-analyzed fMRI data from Holroyd et al. (2018), strictly adhering to the procedures therein (except for the permutation analysis; see below). See Holroyd et al. (2018) for complete details; see also Shahnazian et al. (2022).

#### Participants

Holroyd et al. (2018) included the data of 18 participants (mean age = 23.9yrs; range 19-29yrs; 5 males). Participants had no MRI contra-indications, signed an informed consent form before participating, and were paid for their participation. The experimental protocol was approved by the Ethical Committee of Ghent University Hospital (Belgium) and the University of Victoria Human Research Ethics Board (Canada), and conducted in accordance with the ethical standards described in the 1964 Declaration of Helsinki.

#### Procedure

Participants were first instructed about the rules of the coffee-tea task and then performed 6 practice trials. The instructions emphasized that they should choose to prepare coffee or tea on each trial at random, as if flipping a coin, and similarly to choose the second ingredient (water or cream for coffee; water or sugar for tea) randomly. They completed in total 72 trials, split over four runs of 18 trials each. Between each run, participants could take a short self-paced break. At the end of each run, participants received feedback about the number of prepared coffees and teas.

#### fMRI data acquisition

fMRI images were collected using a Siemens 3T Magnetom Trio Scanner with a 32-channel head coil. At the beginning of each session, high-resolution structural T1-weighted images were acquired using an MP-RAGE. Functional scans were acquired using an echo-planar imaging (EPI) pulse sequence (TR=2000 ms; TE=30 ms; 3 x 3 x 3 mm voxels; flip angle=80°; 33 slices per volume; FOV=224 mm; interleaved slice acquisition) in four separate runs. The first 5 volumes of each run were discarded to allow for T1 equilibrium.

#### fMRI preprocessing

Imaging data were pre-processed and analyzed using the Matlab toolbox SPM12 (Wellcome Trust Centre for Neuroimaging). Images of each participant were re-aligned to the first image in each time series, slice-time corrected using the first slice as reference, co-registered to the mean functional image, and then to the individual anatomical volume.

The General Linear Model (GLM) included 24 regressors of interest, including one for each unique action step in the task at the moment a response was given. They were convolved with the canonical hemodynamic response function (HRF), while the 6 motion parameters and global signal were entered as nuisance regressors. The fMRI time series were high-pass filtered (cut-off 128s) and a first-order autoregressive model was used to correct for temporal autocorrelation.

#### Representational Similarity Analysis (RSA)

In order to identify second-order similarities between how the brain and the RNNs represent the coffee-tea task, we conducted an RSA using a searchlight approach (Kriegeskorte et al., 2008). In particular, we investigated the representational similarity between activity patterns in the brain versus activity patterns in the hidden layer of the three different RNNs described in section 2.2. We expected that the representations associated with the action RDM would correspond most closely to those in caudal ACC, whereas the representations of the goal RDM would correspond most closely to those in rostral ACC.

To facilitate direct comparisons between the two, the brain RDMs were organized the same way as the RNN RDMs. For the brain, we created a 27-element vector of BOLD signal activations, derived from a 3x3x3 voxel cube surrounding a ‘center’ voxel. This was done for each unique step in the task, such that 24 activity patterns were extracted per person per voxel cube. Each 27-element vector thus reflects the activity patterns during one unique action step in the task. Then, for each voxel cube an RDM was constructed by computing the 1 - Spearman rank-order correlation between the activity patterns across all pairwise combinations of the 24 action steps (i.e., a 24x24 matrix). We then vectorized the upper triangle of each RDM, excluding elements along the diagonal, which yielded a 276-unit vector that was used for each ‘center’ voxel in the volume. The 3x3x3 voxel cube was then being moved voxel by voxel through the brain volume, thus ultimately covering the whole brain

Secondly, the 4 RNN-RDMs (see above) and the brain-related RDMs were correlated with each other using a searchlight approach (Kriegeskorte et al., 2008). For each participant, we computed the Spearman’s rank-order correlation between the 276-unit vector of one of the RNNs versus those of the brain per center voxel, such that a correlation value was returned for each voxel in the brain volume. Each correlation thus reflects how similar the activity patterns in and around this voxel were to that of the tested RNN. Next, correlation-values were z-transformed within participants, such that larger values indicated stronger correlations to the RNN compared to other voxels in the brain (Holroyd et al., 2018). Furthermore, data were normalized into MNI space. Lastly, a simple t-test was run across participants per voxel cube, to identify which searchlight volumes exhibited significant second-order similarities with the activation states of the RNN hidden units. Results were family-wise error (FWE) cluster-corrected, using a primary voxel-wise threshold of p < .001.

Note that the original paper also included a permutation analysis that protected against Type-I errors in order to evaluate the strong hypothesis that ACC, as compared to all other regions of the brain, would correlate most strongly with the model RDM (Holroyd et al., 2018). By contrast, here we explored the weaker hypothesis that the inclusion of goal units would reveal a cluster of activity in ACC that was rostral to the cluster of ACC activity observed for the flat network, without predicting their relative sizes. For this reason, we report the largest 3 clusters yielded per RNN.

## Results

Unsurprisingly, our analysis using the Elman network replicated the results of Holroyd et al. (2018). Inspection of the neural network RDM (Figure 10A) reveals that, for all possible sequences, the actions resemble their sequential counterparts across the 4 sequences (for example action 3 of sequence 1 resembles action 3 of sequence 2), yielding a diagonal pattern (blue strips along the diagonal of each sequence). Likewise, the “stir” actions (actions 3 and 5 of all sequences) elicit similar representations, both within and across sequences, producing checkered patterns. Finally, there appears to be an ordering effect such that representations are most similar for states that are closest within a sequence, causing the top right and bottom left of each 6x6 square to be more dissimilar (yellow) than cells closer to the diagonal. Further, when comparing the neural network RDM to the fMRI data, the largest identified cluster corresponded to caudal ACC (Table 4, Figure 10C, yellow) with MNI coordinates and a peak t-value that were identical to those found by Holroyd et al. (2018) (see their Figure 4a).^3^

**Figure 10.**
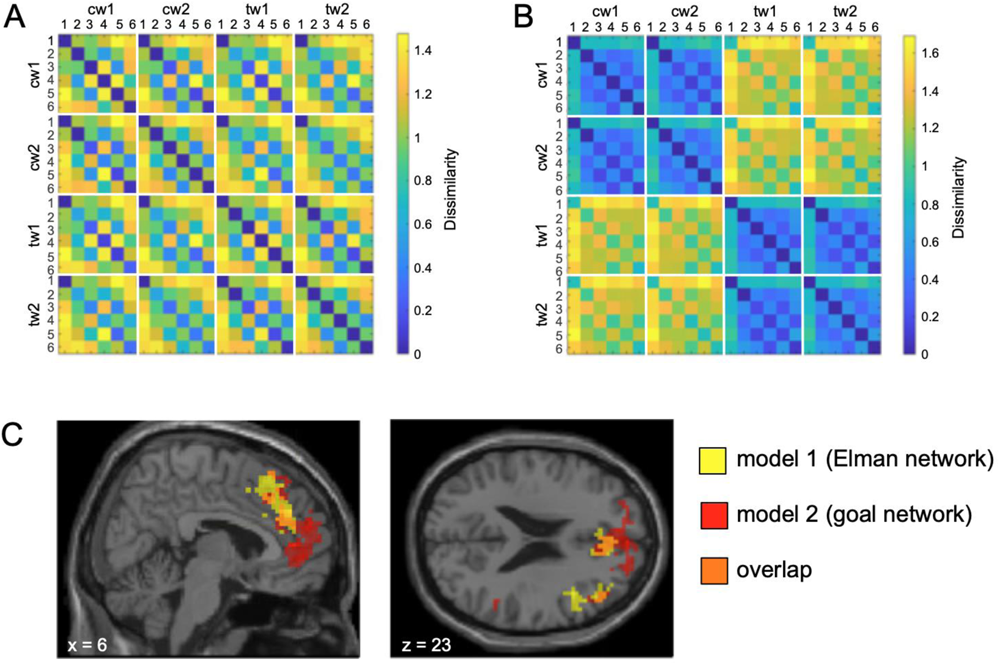
RDMs of the context layer of (A) the Elman network (model 1) and (B) the goal network (model 2). Each row and column of the RDM corresponds to one step of one sequence; for instance the top-right square denotes the distance between step 1 of sequence cw1 (coffee, water first) and step 6 of sequence tw2 (tea, water second). Compared to the Elman network, the RDM of the goal network shows stronger similarity among the separate coffee steps and the separate tea steps (as indicated by the blue color), while the coffee and tea steps are more dissimilar to each other (as indicated by the yellow color). (C) Results of the searchlight analysis for both models, including their overlap, visualized using xjView toolbox (https://www.alivelearn.net/xjview). Compared to the Elman network, the inclusion of goal units yields a larger cluster that includes more anterior regions such as rostral ACC, in line with the idea of a rostral-caudal abstraction gradient. Coordinates are given in MNI space, and clusters are displayed using an uncorrected p-value of .001.

**Table 4:**
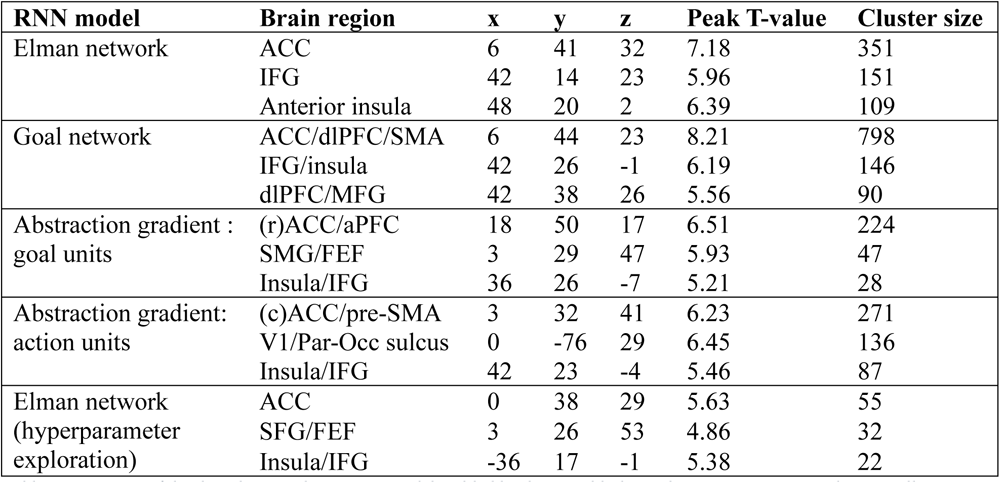
Overview of the three largest clusters per model yielded by the searchlight analysis, using MNI coordinates. All clusters survived family-wise error (FWE) cluster-correction (pFWE=.05) using a primary voxel-wise threshold of p=0.001. ACC=anterior cingulate cortex, IFG = inferior frontal gyrus, dlPFC = dorsolateral prefrontal cortex; SMA = supplementary motor cortex, MFG = middle frontal gyrus, SMG=superior middle gyrus, FEF =frontal eye fields, V1 = primary visual cortex, SFG = superior frontal gyrus.

| RNN model | Brain region | x | y | z | Peak T-value | Cluster size |
| --- | --- | --- | --- | --- | --- | --- |
| Elman network | ACC | 6 | 41 | 32 | 7.18 | 351 |
|  | IFG | 42 | 14 | 23 | 5.96 | 151 |
|  | Anterior insula | 48 | 20 | 2 | 6.39 | 109 |
| Goal network | ACC/dlPFC/SMA | 6 | 44 | 23 | 8.21 | 798 |
|  | IFG/insula | 42 | 26 | -1 | 6.19 | 146 |
|  | dlPFC/MFG | 42 | 38 | 26 | 5.56 | 90 |
| Abstraction gradient :<br>goal units | (r)ACC/aPFC | 18 | 50 | 17 | 6.51 | 224 |
|  | SMG/FEF | 3 | 29 | 47 | 5.93 | 47 |
|  | Insula/IFG | 36 | 26 | -7 | 5.21 | 28 |
| Abstraction gradient:<br>action units | (c)ACC/pre-SMA | 3 | 32 | 41 | 6.23 | 271 |
|  | V1/Par-Occ sulcus | 0 | -76 | 29 | 6.45 | 136 |
|  | Insula/IFG | 42 | 23 | -4 | 5.46 | 87 |
| Elman network<br>(hyperparameter<br>exploration) | ACC | 0 | 38 | 29 | 5.63 | 55 |
|  | SFG/FEF | 3 | 26 | 53 | 4.86 | 32 |
|  | Insula/IFG | -36 | 17 | -1 | 5.38 | 22 |

Figure 10B shows the RDM for the Goal network. To compare the Goal RDM with the Elman RDM, we applied MDS to these distances separately for the two networks (Figure 11). As can be seen by inspection, the incorporation of goal units corresponding to “tea” and “coffee” in the Goal model substantially altered the hidden layer representations: in addition to the patterns observed for the Elman network, the representations are dominated by two separate manifolds within the state space corresponding to the tea and coffee sequences. The addition of goals separates the sequence trajectories into two different clusters, one comprising steps of the tea sequences and the other comprising steps of the coffee sequences. Further, these distinctions are reflected in brain activity: The largest cluster of brain activations showing second-order similarity to the Goal RDM was again located in the ACC, and comprised more voxels than the cluster associated with the Elman network (i.e., 798 voxels; Table 4, Figure 10C). Crucially, as Figure 10C indicates, the cluster identified by the Goal network showed partial overlap with that of the Elman network but extended more rostrally to ventral and rostral ACC, in line with the prediction that ACC would be hierarchically organized along a rostral-caudal abstraction gradient (Holroyd & McClure, 2015).

**Figure 11.**
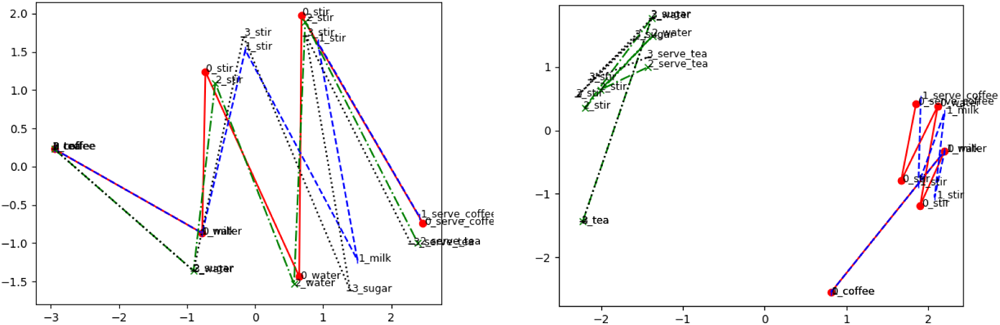
Multi-dimensional scaling of the hidden layer activations of the network at each step of the sequences. The introduction of goal units causes goal representations to become separate; note however that a symmetry between the tea and coffee sequences remains present.

As discussed above, a third network, termed the abstraction gradient network, was trained with a loss function that penalized long-distance connections (simulating wiring costs), inducing a gradient of abstraction in the hidden layer. Thus, in a single network, the hidden units that processed action-related information were prone to communicate with other such units on the same side of the layer, whereas the hidden units that processed goal-related information were prone to communicate with other such units at the other side of the layer. Accordingly, two RDMs were produced, one corresponding to the units associated with the action-focused half of the layer, and one corresponding to the units associated with the goal-focused half of the layer (Figure 12A and 12B). Both RDMs reveal patterns at both the action and goal levels, but with a clear preference depending on the hierarchical level. As predicted, the largest identified cluster was found in the ACC for both the action and goal units (Table 4). Further, the action units targeted the caudal regions more strongly whereas the goal units targeted the rostral regions more strongly (including rostral ACC) (Figure 12C), indicating presence of a rostral-caudal abstraction gradient that mirrors the hierarchical representations encoded within the RNN model.

**Figure 12.**
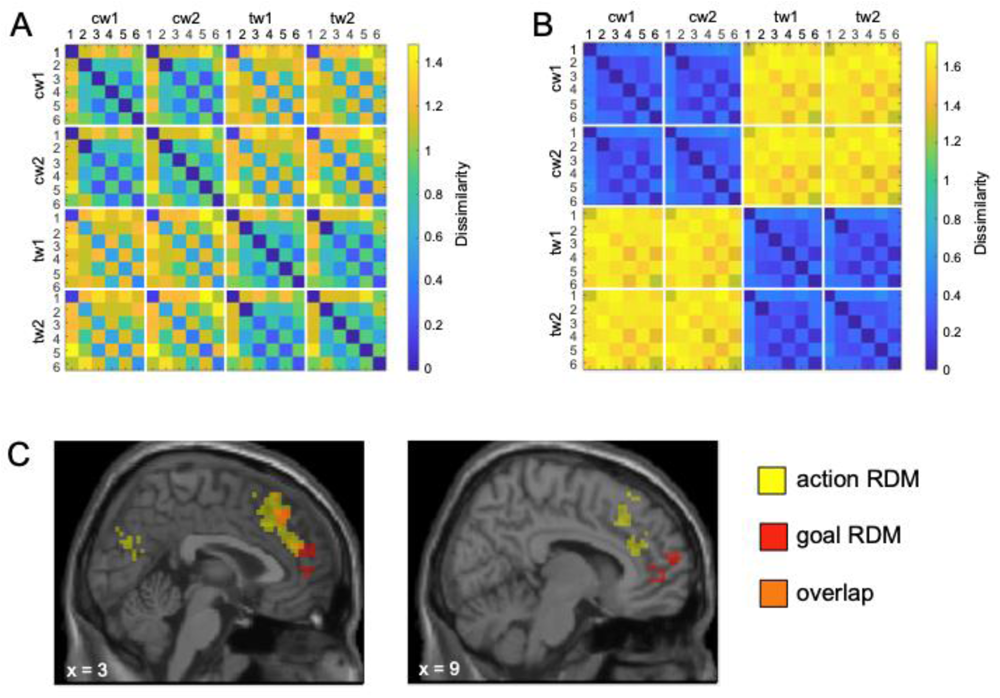
RDMs of the context layer using (A) the ’action’ units and (B) the ’goal’ units from the abstraction gradient model. The RDM based on the goal units clearly separates between coffee and tea, but this separation is still visible in the RDM based on the action units to a lesser extent. (C) Results of the searchlight analysis for the abstraction gradient model, visualized using xjView toolbox (https://www.alivelearn.net/xjview). The rostral-caudal abstraction gradient is visible, such that actions are represented more strongly in the caudal part of the ACC, while goals are represented more strongly in the rostral part of the ACC. Coordinates are given in MNI space, and clusters are displayed using an uncorrected p-value of .001.

Finally, the Elman network hyperparameter exploration RDM confirmed the robustness of the other findings: it detected smaller clusters, as expected, but the top three clusters corresponded to areas found by the other models, with ACC still the largest and most significant cluster.

### Study 2 summary

The use of RSA revealed that ACC contained the largest cluster of voxels whose representations matched those of the Elman network, showing ACC represents temporally extended tasks.This replicated work by Holroyd et al. (2018). Adding goal units to the network led to representations that clearly distinguished tea and coffee sequences: compared to the Elman network, the Goal network representations corresponded to a larger cluster of ACC voxels extending into rostral ACC. This suggests a rostro-caudal gradient in ACC from abstract, temporally extended task features to shorter-term representations more related to individual actions. To confirm this, a third network was constructed that uses wiring costs to generate complementary representations at different levels of abstraction along a gradient in its hidden layer. Application of RSA revealed that the goal side and the action side of this network’s hidden layer corresponded most strongly to representations in rostral ACC and caudal ACC, respectively, consistent with the hierarchical hypothesis.

### General discussion

A basic conflict exists between models that make use of discrete, modular representations that allow for hierarchical organization and componential production of complex behaviors (Cooper et al. 2014), versus models that utilize more distributed representations that are often better at generalizing to novel stimuli (Botvinick & Plaut 2004). Distributed representations tend to be unconstrained and non-compartmentalized, forming abstract patterns that are insensitive to superficial differences in task states. In contrast, hierarchy enables flexible behaviors that minimize interference between similar task states by organizing representations into separate modules. It remains unclear whether this apparent opposition requires arbitration by a higher control system.

Actual brains, which exhibit evidence for both hierarchical and distributed representations, have evidently found a solution to this conundrum. Although much of the brain exhibits features of both types of representations, the tension between them is perhaps most acute in frontal cortex (Badre & Nee, 2018). Frontal cortex is believed to implement processes related to cognitive control, a group of cognitive abilities characterized by top-down regulation of bottom-up information flows that is often modeled hierarchically (e.g., Cohen et al., 1990). The present work explores these issues by first developing a computational model that incorporates both hierarchical and distributed properties, and then examining these properties against neural data with a focus on the ACC, a frontal system implicated in hierarchical control over goal-directed action sequences.

We began by studying the behavior of models that combined elements of hierarchy and distributed representations. Hierarchy was achieved by making use of discrete “goal units” corresponding to specific sets of action sequences, and distributed representations were obtained by training a recurrent neural network (via backpropagation through time) on these sequences. Goal units or variants thereof have often been used in artificial neural networks to implement temporally extended behavior in a hierarchically structured manner, both in cognitive models (Cooper et al., 2014) and in AI (Bacon et al., 2017). We expected these methods to reveal characteristics of both hierarchical and distributed systems.

### The computational effects of hierarchy on distributed representations

In study 1, two types of RNN models, one with goal units and one without, were trained within a simulated kitchen environment to produce 21 different coffee or tea-making sequences. This task was characterized by two properties that the networks could exploit. First, the sequences could be decomposed into a small number of subsequences, ensuring that the task could be performed by recombining such subsequences, potentially making a hierarchical representation of the task advantageous. Second, many similarities were present at the action-, sub-sequence-, and sequence-levels, which themodel could encode with distributed representations to enable generalization to novel variations in the task. We reasoned that the goal units would constrain and guide learning, shaping the network to represent higher-level abstract patterns. This in turn would help to maintain stable task performance even when task execution was disturbed, for example, in case of environmental distractors or competition from other tasks. Goal units might be especially helpful in reducing RNN instability, compared to a feedforward network.

In particular, we expected that this scaffolding would cause the network to maintain the correct, temporally extended behavior even when subjected to stress. Further we expected that failures would occur in a more structured way compared to the Elman network, namely by replacing one extended sequence of actions with another rather than by executing wrong actions in isolation. To explore this possibility, we added noise to the activation values of the hidden units in order to simulate interference caused by non-task-related processing. However, contrary to our expectations, we found that the goal units exerted only limited influence over network behavior, as these networks performed about as well as those lacking goal units, and in a qualitatively similar fashion.

This result may be an example of the argument that, given sufficient computational power, flexible methods (such as an unconstrained RNN) tend to outperform ad-hoc architectures (Sutton, 2019). In the present case, the specific hierarchical decomposition of the task provided a hierarchical bias as input to the network, but it was striking that the scaffolding provided by these goals and subgoals had such limited effects on the observed behavior. This result is reminiscent of similar observations in the HRL literature, for instance, that an option-based architecture yielded limited improvements in performance and learning relative to a less complex architecture (Bacon et al., 2017). Nevertheless, that learning algorithm demonstrated a significant performance advantage when required to adapt to a non-stationary environment.^4^

In such environments, when confronted with tasks in which reflexive or automatic behaviors fail, humans may engage higher cognitive abilities such as the application of control. In the modeling literature, Cooper et al. (2014) have proposed models that implement hierarchical structure explicitly for the sake of applying control. Their goal units interface between systems specializing on different abstract levels of decision-making, simulating an executive system that modulates action selection by the lower-level system. Corroborating their results, the control interventions in study 1 showed that modifying the activation of the goal units, after training, could extend classical effects associated with cognitive control (Cohen et al., 1990) to the domain of hierarchical action sequences. In these simulations, the goal units could be used to select a less-preferred action sequence in place of a more-preferred one (such as making coffee with milk when cream was normally preferred), and could also be used to recombine subsequences in order to produce a novel sequence with no additional training (such as making tea with cream). Further, applying more control allowed for overcoming the inertia of habit more reliably.

Collectively, these results generalize classic accounts of the flexibility afforded by cognitive control mechanisms (Cohen, 2017) to the production of hierarchical action sequences. The mechanism by which goal units regulate a lower-level system has received substantial attention, mostly focused on feedforward networks since the seminal work of Cohen, Dunbar, and McClelland (1990). Whereas these systems are comparatively well-understood, we were interested in sequential decision making, which our model learned by making use of recurrent connections to preserve contextual information. In this less-explored setting, control may be applied over time, affecting the representations as they evolve on each time-step. We examined the effect of goal interventions on the representational distance patterns of hidden layer representations.

Musslick and Cohen (2021) have proposed that control serves to “deepen” attractor states (making it more difficult for the agent to exit the attractor basin of the current goal, thereby preventing accidental goal-switching). In representational terms, we expected the application of control to increase the representational dissimilarity – the distance – between the correct state and the conflicting incorrect state, preventing noise or distractors from causing an accidental task switch. Consistent with this interpretation, we found that multiplying the activation values of the goal units by a constant increased the Euclidian distance between states corresponding to different goals compared to states corresponding to the same goal. In our work, this effect emerged without a dedicated mechanism, though it was not a very strong effect. It is possible that the effect would become more pronounced if the networks were trained in a stochastic or noisy environment (which could punish shallower attractors), or if it were trained in environments requiring control-induced variability in behavior, i.e. exploration or play, which might reward reliable goal units.

We next investigated the degree to which real brains implement these hierarchically organized distributed representations. Towards this end, we extended our prior work that used an Elman network to identify distributed representations of sequence production in ACC (Holroyd et al., 2018). Using RSA, we compared predictions of the Elman and Goal models to brain representations of sequence production as obtained in humans using fMRI. The Goal model RSA yielded a larger cluster of ACC voxels relative to the Elman network RSA, suggesting that distributed and hierarchically-organized representations of action sequences coexist within the brain. Furthermore, the cluster associated with the Goal model, which was characterized by a greater degree of higher abstraction, extended more rostrally than the cluster associated with the Elman model, consistent with a rostro-caudal organization of representations in the medial cortex.

However, this apparent hierarchical organization was revealed using two separate models: the Elman model and the Goal model. Better support for this inference would be provided by a single model incorporating both goal-related and action-related units in the hidden layer. Toward this end, and motivated by arguments that processing delays and metabolic costs increase with axon length (Chen et al., 2006), we trained the Gradient model, which incorporated a cost function that penalized long connection lengths, In line with our predictions, the RSA revealed that human ACC is organized similarly to the Gradient model, namely implementing high-vs. low-level representations of task progress along a rostro-caudal gradient. These results suggest that ACC encodes task characteristics using complementarily distributed and hierarchical representations, perhaps explaining some of the diversity of functions associated with ACC (Shahnazian & Holroyd, 2018; Holroyd & Verguts, 2021).

### Interpretation of the gradient and limitations

Study 1 suggested a possible role for a segregation of task representations according to their level of abstraction. In this view, abstract units guide network behavior by recombining abstract policy components to adapt to novel demands, or by facilitating exploratory behaviors. In a machine learning context, Vezhnevets et al. (2017) have shown that a distributed goal representation can act as a “manager” to direct the behavior of a lower-level network. The combination of hierarchy and distributed representations might enable generalization at each level of abstraction, while interactions between the abstraction levels could endow the network with greater flexibility. The findings of study 2, which reveal a gradient of abstraction in the ACC, are consistent with this view.

Nevertheless, this straightforward interpretation must be qualified in light of both technical limitations and the broader evidence regarding ACC function. In particular, the models used here were highly abstract and, as such, simulated aspects of ACC function only to a first-order approximation. The models are shallow neural networks (having only one hidden layer) with a small number of units and were trained using the biologically implausible supervised learning signal (see e.g. Lillicrap et al., 2020). Further, the unit activations were fully updated on each time step, which overly simplifies the temporal dynamics at play in actual neural systems. Such simple models constrain the solution space by offering fewer degrees of freedom to the experimenter, but in so doing elide many of the details of neural implementation.

In general, neural network representations are not well-understood. Recent explorations have demonstrated that they are highly dependent on network architecture and on the choice of hyperparameters (Wang et al., 2018, Maheswaaran et al., 2019, Storrs et al., 2021, Flesch et al., 2022). By considering second order representational similarities, rather than direct correspondences, RSA can circumvent some of these complexities (Lu et al., 2019). For example, recent successes using RSA suggest that, despite the many large differences with their biological counterparts, artificial neural networks do indeed yield second-order similarities with neural data that are robust to differences in network parameterization (e.g., Cichy et al., 2016, Bakhtiari et al., 2021). To address some of these concerns, we trained multiple Elman networks across a vast range of hyperparameters combinations and observed that the average of the RDMs across these networks also identified an ACC cluster. Additionally, all our RSAs (Elman, goal, and abstraction gradient networks) were based on an average RDM from one hundred network instances, likely minimizing variance in the results, compared to studies which use a small number of larger networks – often just one.

Another consideration concerns not the specific choices of implementation but rather the proposed function of the network. Crucially, the model simulates the patterns of activity within ACC during the production of action sequences but it is not intended to instantiate ACC function itself. The RSA technique can reveal similarity between representations, but that is not sufficient to establish conclusively identity of function. Although the representations in the hidden layer of the neural network models are used to support action and/or goal selection, the ACC appears to use similar representations for something more subtle. Early debates about ACC function hinged on the question of whether ACC contributed directly to action production (an “actor”), or whether it monitored task performance for control purposes (a “monitor”) (Holroyd & Yeung, 2003). However, it seems ACC has only a weak modulatory influence over human behavior, as ACC lesions tend to disrupt normal behavior only minimally (Holroyd & Yeung, 2012). Rather, we expect that ACC representations reflect a role in monitoring the state of the system during task progression and are used to modulate the activity of other neural systems that are more directly responsible for producing behavior, for example lateral prefrontal cortex (Aarts et al., 2008, Holroyd & Yeung, 2012, Holroyd, 2023).

Future work should incorporate both actor and monitor components in a single simulation, directly specificying the supporting role of ACC for the activities of other neurocognitive components of the agent. This role could entail extracting abstract, temporally extended task features—such as preserving the task context, predicting outcomes, or assessing expected values—consistent with our previous work (Holroyd et al., 2018) and with other recent models of ACC (Akam et al., 2021, Alexander & Brown, 2021, and others; see Vassena et al., 2017, for a review). Perhaps this variety of representations is useful to brain networks serving different purposes, contributing to the illusion that ACC does “everything”. Our work here suggests that ACC not only keeps track of the hidden state of the world, but also represents the agent’s own behavior at different levels of abstraction, an organization that furthers control over the execution of high-level action sequences.

## Author contributions

Thomas R. Colin: Software, writing (original draft). Iris Ikink: Formal analysis, writing (review and editing). Clay Holroyd: Supervision, writing (review and editing).

## Data availability statement

fMRI data has been deposited in the Open Science Framework repository (https://osf.io/UHJCF/; https://doi.org/10.17605/OSF.IO/WXHTA). Contact the corresponding author for computational modeling details and code.

## Footnotes

1 The term “ablation studies” is used here in the artificial intelligence sense, where the approach is used to test the role of the different components of the model. It is not intended to simulate actual brain lesions.

2 A one-hot vector has exactly one entry set to 1 and the others set to 0.

3 The ACC cluster consisted of 351 voxels instead of 348 as reported by Holroyd et al. (2018). We suspect that this discrepancy results from differences between analysis software used in these two studies (i.e., AFNI vs. SPM12).

4 Others, such as Vezhnevets et al. (2017), observed performance improvements using an ingenuous algorithm making use of distributed goal representations; however, they view these improvements as resulting mainly from long-range credit assignment and exploration, issues that were not present in the same form in our model due to the supervised training scheme. McGovern and Sutton (1998) offer an accessible discussion on the benefits of macro-actions in an RL setting.

## Appendix List of training sequences: study 1

### Subsequences

**Table 5:** Subsequences in the kitchen environment.

|  |  |
| --- | --- |
| **Add cream** | Fixate fridge; open (if no cream is present, switch to “add milk” from step 3, else continue); fixate cream; take; fixate mug; add to mug; fixate fridge; put down; close |
| **Add grounds** | Fixate cupboard; open; fixate coffee jar; take; open; fixate mug; add to mug; fixate coffee jar; close; fixate cupboard; put down |
| **Add milk** | Fixate fridge; open; fixate milk; take; fixate mug; add to mug; fixate fridge; put down; close |
| **Add sugar** | Fixate sugar box; take; fixate mug; add to mug |
| **Dip teabag** | Fixate cupboard; open; fixate teabags; take; fixate mug; add to mug |
| **Drink coffee** | Fixate mug; take; sip; fixate table; put down; say “good coffee” |
| **Drink tea** | Fixate mug; take; sip; fixate table; put down; say “good tea” |
| **Tidy up** | (if not already being fixed) fixate cupboard; close |
| **Stir** | Fixate spoon; take; fixate mug; stir; fixate table; put down |

### Sequences

**Table 6:** Sequences in the kitchen environment.

| Sequence | Environment | Instructions | Subsequences |
| --- | --- | --- | --- |
| Coffee |  | Coffee | Add grounds; tidy up; stir; drink coffee |
| Coffee sugar |  | Coffee, sugar | Add grounds; stir; add sugar; tidy up; drink coffee |
| Coffee 2 sugars |  | Coffee, sugar, extra sugar | Add grounds, stir; add sugar; stir; add sugar; tidy up; stir; drink coffee |
| Coffee cream | Cream present | Coffee, dairy | Add grounds; tidy up; stir; add cream; stir; drink coffee |
| Coffee cream sugar | Cream present | Coffee, dairy, sugar | Add grounds; stir; add cream; stir; add sugar; tidy up; stir; drink coffee |
| Coffee<br>cream 2<br>sugars | Cream<br>present | Coffee,<br>dairy, sugar,<br>extra sugar | Add grounds; stir; add cream; stir; add sugar;<br>stir; add sugar; tidy up; stir; drink coffee |
| Coffee<br>sugar<br>cream | Cream<br>present | Coffee,<br>dairy, sugar | Add grounds; stir; add sugar; tidy up; stir; add<br>cream; stir; drink coffee |
| Coffee 2<br>sugars<br>cream | Cream<br>present | Coffee,<br>dairy, sugar,<br>extra sugar | Add grounds; stir; add sugar; stir; add sugar;<br>tidy up; stir; add cream; stir; drink coffee |
| Coffee<br>milk | Cream absent | Coffee,<br>dairy | Add grounds; tidy up; stir; add cream (find out<br>there's no cream); add milk; stir; drink coffee |
| Coffee<br>milk<br>sugar | Cream absent | Coffee,<br>dairy, sugar | Add grounds; stir; add cream (find out there's<br>no cream); add milk; stir; add sugar; tidy up;<br>stir; drink coffee |
| Coffee<br>sugar<br>milk | Cream absent | Coffee,<br>dairy, sugar,<br>extra sugar | Add grounds; stir; add sugar; tidy up; stir; add<br>cream (find out there's no cream); add milk;<br>stir; drink coffee |
| Coffee<br>milk 2<br>sugars | Cream absent | Coffee,<br>dairy, sugar | Add grounds; stir; add cream (find out there's<br>no cream); add milk; stir; add sugar; stir; add<br>sugar; tidy up; stir; drink coffee |
| Coffee 2<br>sugars<br>milk | Cream absent | Coffee,<br>dairy, sugar,<br>extra sugar | Add grounds; stir; add sugar; stir; add sugar;<br>tidy up; stir; add cream (find out there's no<br>cream); add milk; stir; drink coffee |
| Tea |  | Tea | Dip teabag; tidy up; stir; drink tea |
| Tea sugar |  | Tea, sugar | Dip teabag; stir; add sugar; tidy up; stir; drink<br>tea |
| Tea 2<br>sugars |  | Tea, sugar,<br>extra sugar | Dip teabag; stir; add sugar; stir; add sugar; tidy<br>up; stir; drink tea |
| Tea milk |  | Tea, dairy | Dip teabag; tidy up; stir; add milk; stir; drink<br>tea |
| Tea milk<br>sugar |  | Tea, dairy,<br>sugar | Dip teabag; stir; add milk; stir; add sugar; tidy<br>up; stir; drink tea |
| Tea sugar<br>milk |  | Tea, dairy,<br>sugar | Dip teabag; stir; add sugar; tidy up; stir; add<br>milk; stir; drink tea |
| Tea milk<br>2 sugars |  | Tea, dairy,<br>sugar, extra<br>sugar | Dip teabag; stir; add milk; stir; add sugar; stir;<br>add sugar; tidy up; stir; drink tea |
| Tea 2<br>sugars<br>milk |  | Tea, dairy,<br>sugar, extra<br>sugar | Dip teabag; stir; add sugar; stir; add sugar; tidy<br>up; stir; add milk; stir; drink tea |

### No effect of goal units on dimensionality

Flesch and colleagues (2022) have argued that lower dimensionality is associated with higher generalizability, enabling novel states to be processed according to learned state-action mappings; in contrast, higher dimensionality creates affordances for new stimulus-action pairings overruling learned behavior, which might be leveraged by a higher-level control system.

The dimensionality of representations was calculated by evaluating the rank of the activation matrix of each network over all sequences and steps (see Badre et al., 2021) for a discussion on dimensionality measurement). This analysis did not reveal any significant changes in dimensionality across standard and increased (x2) control levels, evaluated using a tolerance of 0.05 (corresponding to the noise magnitude at which few errors occur). We investigated whether other tolerance levels showed different results, and found no significant effect for sampling tolerance levels ranging from 0.01 (where matrix rank is 50 for most networks) to 150 (where it matrix rank is 1 for most networks), confirming that goal units had no effects on representational dimensionality, as measured based on rank.

This negative result suggests that such dimensionality differences do not emerge from supervised training via backpropagation. It remains possible that other training methods allow them to appear in an emergent fashion. Failing that, a dedicated mechanism could be required.

## References

1. Aarts, E., Roelofs, A., & van Turennout, M. (2008). Anticipatory activity in anterior cingulate cortex can be independent of conflict and error likelihood. The Journal of Neuroscience : the Official Journal of the Society for Neuroscience, 28(18), 4671–4678. 10.1523/JNEUROSCI.4400-07.2008

2. Akam, T., Rodrigues-Vaz, I., Marcelo, I., Zhang, X., Pereira, M., Oliveira, R. F., Dayan, P., & Costa, R. M. (2021). The anterior cingulate cortex predicts future states to mediate model-based action selection. Neuron, 109(1), 149–163.e7. 10.1016/j.neuron.2020.10.013

3. Alexander, W. H., & Brown, J. W. (2011). Medial prefrontal cortex as an action-outcome predictor. Nature Neuroscience, 14(10), 1338–1344. 10.1038/nn.2921

3a. Alexander, W. H., & Womelsdorf, T. (2021). Interactions of medial and lateral prefrontal cortex in hierarchical predictive coding. Frontiers in Computational Neuroscience, 15, 605271. 10.3389/fncom.2021.605271

4. Bacon, P.-L., Harb, J., & Precup, D. (2017). The option-critic architecture. Proceedings of the AAAI Conference on Artificial Intelligence, 31(1). 10.1609/aaai.v31i1.10916

5a. Badre, D., Bhandari, A., Keglovits, H., & Kikumoto, A. (2021). The dimensionality of neural representations for control. Current Opinion in Behavioral Sciences, 38, 20–28. 10.1016/j.cobeha.2020.07.002

5. Badre, D., & Desrochers, T. M. (2019). Hierarchical cognitive control and the frontal lobes. Handbook of Clinical Neurology, 163, 165–177. 10.1016/B978-0-12-804281-6.00009-4

6. Badre, D., & Nee, D. E. (2018). Frontal cortex and the hierarchical control of behavior. Trends in Cognitive Sciences, 22(2), 170–188. 10.1016/j.tics.2017.11.005

7. Bakhtiari, S., Mineault, P., Lillicrap, T., Pack, C., & Richards, B. (2021). The functional specialization of visual cortex emerges from training parallel pathways with self-supervised predictive learning. Advances in Neural Information Processing Systems, 34.

8. Botvinick, M. M., & Bylsma, L. M. (2005). Distraction and action slips in an everyday task: evidence for a dynamic representation of task context. Psychonomic Bulletin & Review, 12(6), 1011–1017. 10.3758/bf03206436

9. Botvinick, M., & Plaut, D. C. (2004). Doing without schema hierarchies: a recurrent connectionist approach to normal and impaired routine sequential action. Psychological Review, 111(2), 395–429. 10.1037/0033-295X.111.2.395

10. Bowers J. S. (2017). Parallel distributed processing theory in the age of deep networks. Trends in Cognitive Sciences, 21(12), 950–961. 10.1016/j.tics.2017.09.013

10a. Bullmore, E., & Sporns, O. (2012). The economy of brain network organization. Nature Reviews Neuroscience, 13(5), 336–349. 10.1038/nrn3214

11. Chen, B. L., Hall, D. H., & Chklovskii, D. B. (2006). Wiring optimization can relate neuronal structure and function. Proceedings of the National Academy of Sciences of the United States of America, 103(12), 4723–4728. 10.1073/pnas.0506806103

12. Cherniak C. (1994). Component placement optimization in the brain. The Journal of Neuroscience, 14(4), 2418–2427. 10.1523/JNEUROSCI.14-04-02418.1994

12a. Chklovskii, D. B., & Koulakov, A. A. (2004). Maps in the brain: what can we learn from them?. Annual Review of Neuroscience, 27, 369–392. 10.1146/annurev.neuro.27.070203.144226

13. Chomsky, N. (1957), Logical structures in language. American Documentation, 8 : 284–291. https ://doi.org/10.1002/asi.5090080406

14. Christoff, K., Keramatian, K., Gordon, A. M., Smith, R., & Mädler, B. (2009). Prefrontal organization of cognitive control according to levels of abstraction. Brain Research, 1286, 94–105. 10.1016/j.brainres.2009.05.096

15. Cichy, R. M., Khosla, A., Pantazis, D., Torralba, A., & Oliva, A. (2016). Comparison of deep neural networks to spatio-temporal cortical dynamics of human visual object recognition reveals hierarchical correspondence. Scientific Reports, 6, 27755. 10.1038/srep27755

16. Cohen, J. D. (2017). Cognitive control: Core constructs and current considerations. The Wiley handbook of cognitive control, 1-28. 10.1002/9781118920497.ch1

17. Cohen, J. D., Dunbar, K., & McClelland, J. L. (1990). On the control of automatic processes: A parallel distributed processing account of the Stroop effect. Psychological Review, 97(3), 332–361. 10.1037/0033-295x.97.3.332

18. Cooper, R. P., Ruh, N., & Mareschal, D. (2014). The goal circuit model: A hierarchical multi-route model of the acquisition and control of routine sequential action in humans. Cognitive Science, 38(2), 244–274. 10.1111/cogs.12067

19. Flesch, T., Juechems, K., Dumbalska, T., Saxe, A., & Summerfield, C. (2022). Orthogonal representations for robust context-dependent task performance in brains and neural networks. Neuron, 110(7), 1258–1270.e11. 10.1016/j.neuron.2022.01.005

20. Gardner, H. (1985). The mind’s new science: A history of the cognitive revolution. Basic Books.

21. Garnelo, M. & Shanahan, M. (2019). Reconciling deep learning with symbolic artificial intelligence: representing objects and relations. Current Opinion in Behavioral Sciences, 29, 17–23. 10.1016/j.cobeha.2018.12.010.

22. Grafman J. (1995). Similarities and distinctions among current models of prefrontal cortical functions. Annals of the New York Academy of Sciences, 769, 337–368. 10.1111/j.1749-6632.1995.tb38149.x

23. Haarnoja, T., Hartikainen, K., Abbeel, P., & Levine, S. (2018). Latent space policies for hierarchical reinforcement learning. Proceedings of the 35^th^ International Conference on Machine Learning, PMLR 80:1851-1860.

24. Hanson, S., & Burr, D. (1990). What connectionist models learn: Learning and representation in connectionist networks. Behavioral and Brain Sciences, 13(3), 471–489. 10.1017/S0140525X00079760

25. He, K., Zhang, X., Ren, S., & Sun, J. (2015). Delving deep into rectifiers: Surpassing human-level performance on ImageNet classification. IEEE International Conference on Computer Vision, 1502. 10.1109/ICCV.2015.123.

26. Holroyd, C. (2023). The Controllosphere: The neural origin of cognitive effort. 10.31234/osf.io/2qmhg

27. Holroyd, C. B., & McClure, S. M. (2015). Hierarchical control over effortful behavior by rodent medial frontal cortex: A computational model. Psychological Review, 122(1), 54–83. 10.1037/a0038339

28. Holroyd, C. B., Ribas-Fernandes, J. J. F., Shahnazian, D., Silvetti, M., & Verguts, T. (2018). Human midcingulate cortex encodes distributed representations of task progress. Proceedings of the National Academy of Sciences of the United States of America, 115(25), 6398–6403. 10.1073/pnas.1803650115

29. Holroyd, C. B., & Verguts, T. (2021). The best laid plans: computational principles of anterior cingulate cortex. Trends in Cognitive Sciences, 25(4), 316–329. 10.1016/j.tics.2021.01.008

30. Holroyd, C. B., & Yeung, N. (2003). Alcohol and error processing. Trends in Neurosciences, 26(8), 402–404. 10.1016/S0166-2236(03)00175-9

31. Holroyd, C. B., & Yeung, N. (2012). Motivation of extended behaviors by anterior cingulate cortex. Trends in Cognitive Sciences, 16(2), 122–128. 10.1016/j.tics.2011.12.008

32. Kingma, D. & Ba, J. (2015) Adam: A method for stochastic optimization. *Proceedings of the 3^rd^ International Conference on Learning Representations*.

33. Koechlin, E., Ody, C., & Kouneiher, F. (2003). The architecture of cognitive control in the human prefrontal cortex. Science, 302(5648), 1181–1185. https ://doi.org/10.1126/science.1088545

34. Kriegeskorte, N., Mur, M., & Bandettini, P. (2008). Representational similarity analysis – connecting the branches of systems neuroscience. Frontiers in Systems Neuroscience, 2, 4. 10.3389/neuro.06.004.2008

35. Lake, B. M., Ullman, T. D., Tenenbaum, J. B., & Gershman, S. J. (2017). Building machines that learn and think like people. Behavioral and Brain Sciences, 40, e253. 10.1017/S0140525X16001837

36. Lashley, K. S. (1951). The problem of serial order in behavior. L.A. Jeffress (Ed.), Cerebral mechanisms in behavior, Wiley, New York (1951), pp. 112-131.

37. Lillicrap, T. P., Santoro, A., Marris, L., Akerman, C. J., & Hinton, G. (2020). Backpropagation and the brain. Nature Reviews Neuroscience, 21(6), 335–346. 10.1038/s41583-020-0277-3

38. Lu, Q., Chen, P. H., Pillow, J. W., Ramadge, P. J., Norman, K. A., & Hasson, U. (2018). Shared representational geometry across neural networks. *Integration of Deep Learning Theories workshop*, NeurIPS 2018.

39. McGovern, A., & Sutton, R. S. (1998). Macro-actions in reinforcement learning: an empirical analysis. Computer Science Department Faculty Publication Series.

40. Marblestone, A. H., Wayne, G., & Kording, K. P. (2016). Toward an integration of deep learning and neuroscience. Frontiers in Computational Neuroscience, 10, 94. 10.3389/fncom.2016.00094

41. Miller, E. K., & Cohen, J. D. (2001). An integrative theory of prefrontal cortex function. Annual Review of Neuroscience, 24, 167–202. 10.1146/annurev.neuro.24.1.167

41a. Miller, G. A., Galanter, E., & Pribram, K. H. (1960). Plans and the Structure of Behavior. New York: Holt.

42. Musslick, S., & Cohen, J. D. (2021). Rationalizing constraints on the capacity for cognitive control. Trends in Cognitive Sciences, 25(9), 757–775. 10.1016/j.tics.2021.06.001

43. Richards, B. A., Lillicrap, T. P., Beaudoin, P., Bengio, Y., Bogacz, R., Christensen, A., Clopath, C., Costa, R. P., de Berker, A., Ganguli, S., Gillon, C. J., Hafner, D., Kepecs, A., Kriegeskorte, N., Latham, P., Lindsay, G. W., Miller, K. D., Naud, R., Pack, C. C., Poirazi, P., Schapiro, A. C., Senn, W., Wayne, G., Yamins, D., Zenke, F., Zylberberg, J., Therien, D., Kording, K. P. (2019). A deep learning framework for neuroscience. Nature Neuroscience, 22(11), 1761–1770. 10.1038/s41593-019-0520-2

44. Rogers, T. T., & McClelland, J. L. (2014). Parallel distributed processing at 25: Further explorations in the microstructure of cognition. Cognitive Science, 38(6), 1024–1077. 10.1111/cogs.12148

45. Schank, R. C., & Abelson, R. P. (1977). Scripts, plans, goals and understanding: An inquiry into human knowledge structures. Lawrence Erlbaum.

46. Schmidhuber J. (2015). Deep learning in neural networks: an overview. Neural Networks: the Official Journal of the International Neural Network Society, 61, 85–117. 10.1016/j.neunet.2014.09.003

47. Shahnazian, D., & Holroyd, C. B. (2018). Distributed representations of action sequences in anterior cingulate cortex: A recurrent neural network approach. Psychonomic Bulletin & Review, 25(1), 302–321. 10.3758/s13423-017-1280-1

48. Shahnazian, D., Senoussi, M., Krebs, R. M., Verguts, T., & Holroyd, C. B. (2022). Neural representations of task context and temporal order during action sequence execution. Topics in Cognitive Science, 14(2), 223–240. 10.1111/tops.12533

49. Storrs, K. R., Kietzmann, T. C., Walther, A., Mehrer, J., & Kriegeskorte, N. (2021). Diverse deep neural networks all predict human inferior temporal cortex well, after training and fitting. Journal of Cognitive Neuroscience, 33(10), 2044–2064. 10.1162/jocn_a_01755

50. Sutton, R. S. (2019). The bitter lesson. http://www.incompleteideas.net/IncIdeas/BitterLesson.html

51. Sutton, R.S., Precup, D., & Singh, S. (1999). Between MDPs and Semi-MDPs: A Framework for Temporal Abstraction in Reinforcement Learning. Artificial Intelligence, 112, 181-211. 10.1016/s0004-3702(99)00052-1

52. Van der Maaten, L. & Hinton, G. (2008). Visualizing data using t-SNE. Journal of Machine Learning Research, 9(*86*), 2579–2605.

53. Vassena, E., Holroyd, C. B., & Alexander, W. H. (2017). Computational Models of Anterior Cingulate Cortex: At the Crossroads between Prediction and Effort. Frontiers in neuroscience, 11, 316. 10.3389/fnins.2017.00316

54. Wang, L., Hu, L., Gu, J., Hu, Z., Wu, Y., He, K., & Hopcroft, J. (2018). Towards understanding learning representations: To what extent do different neural networks learn the same representation. Advances in Neural Information Processing Systems, 31.

